# Synaptic integration of subquantal neurotransmission by co-localized G protein coupled receptors in presynaptic terminals

**DOI:** 10.1101/2020.11.12.362491

**Authors:** Emily Church, Edaeni Hamid, Zack Zurawski, Mariana Potcoava, Eden Flores-Barrera, Adriana Caballero, Kuei Y. Tseng, Simon Alford

**Author notes:** These authors contributed equally to this work (Emily Church, Edaeni Hamid). Corresponding Author – Simon Alford.

## Abstract

In presynaptic terminals, membrane-delimited G_i/o_-mediated presynaptic inhibition is ubiquitous and acts through Gβγ to inhibit Ca^2+^ entry, or directly at SNARE complexes to inhibit Ca^2+^-dependent synaptotagmin-SNARE complex interactions. At CA1-subicular presynaptic terminals 5-HT_1B_ and GABA_B_ receptors colocalize. GABA_B_ receptors inhibit Ca^2+^ entry, whereas 5-HT_1B_ receptors target SNARE complexes. We demonstrate in male and female rats that GABA_B_ receptors receptors alter P_r_, whereas 5-HT_1B_ receptors reduce evoked cleft glutamate concentrations allowing differential inhibition of AMPA and NMDA receptor EPSCs. This reduction in cleft glutamate concentration was confirmed by imaging glutamate release using a genetic sensor (iGluSnFR).Simulations of glutamate release and postsynaptic glutamate receptor currents were made. We tested effects of changes in vesicle numbers undergoing fusion at single synapses, relative placement of fusing vesicles and postsynaptic receptors, and the rate of release of glutamate from a fusion pore. Experimental effects of P_r_ changes, consistent with GABA_B_ receptor effects, were straightforwardly represented by changes in numbers of synapses. The effects of 5-HT_1B_ receptor-mediated inhibition are well-fit by simulated modulation of the release rate of glutamate into the cleft. Colocalization of different actions of GPCRs provide synaptic integration within presynaptic terminals. Train-dependent presynaptic Ca^2+^ accumulation forces frequency-dependent recovery of neurotransmission during 5-HT_1B_ receptor activation. This is consistent with competition between Ca^2+^-synaptotagmin and Gβγ at SNARE complexes. Thus, stimulus trains in 5-HT_1B_ receptor agonist unveil dynamic synaptic modulation and a sophisticated hippocampal output filter that itself is modulated by colocalized GABA_B_ receptors which alter presynaptic Ca^2+^. In combination these pathways allow complex presynaptic integration.

**Significance Statement:** Two G protein coupled receptors colocalize at presynaptic sites, to mediate presynaptic modulation by Gβγ, but one – a GABAB receptor inhibits Ca^2+^ entry while another – a 5-HT1B receptor competes with Ca^2+^-synaptotagmin binding to the synaptic vesicle machinery. We have investigated downstream effects of signaling and integrative properties of these receptors. Their effects are profoundly different. GABAB receptors alter Pr leaving synaptic properties unchanged, while 5-HT1B receptors fundamentally change properties of synaptic transmission, modifying AMPA receptor but sparing NMDA receptor responses. Co-activation of these receptors allows synaptic integration because of convergence of GABAB receptor alteration on Ca^2+^ and the effect of this altered Ca^2+^ signal on 5-HT1B receptor signaling. This presynaptic convergence provides a novel form of synaptic integration.

## Introduction

G protein-coupled receptors (GPCRs) control neurotransmitter release at all synapses via membrane delimited actions of Gβγ (Betke et al., 2012). Much attention has been placed on G_i/o_ G protein modulation of Ca^2+^ entry (Tedford and Zamponi, 2006). This alters synaptic vesicle fusion probability (P_r_) (Hessler et al., 1993); a phenomenon taken as proof of presynaptic action (Dobrunz and Stevens, 1997). However, at least one other membrane delimited action of Gβγ occurs. At synapses in lamprey, in amygdala, hippocampus, and in chromaffin and PC12 cells, GPCRs modulate exocytosis via Gβγ-SNARE complex interactions (Blackmer et al., 2001; Takahashi et al., 2001; Blackmer et al., 2005; Chen et al., 2005). This modulation is caused by Gβγ competition with synaptotagmin at c-terminal SNAP-25 on SNARE complexes of primed vesicles. Mutation of this SNAP-25/Gβγ interaction site in mice demonstrates central effects on anxiety, spatial memory, and motor control, as well as endocrine release and metabolism implying broad implications for this exocytotic modulation in brain and the periphery (Zurawski et al., 2019). The mechanistic effects of these different modulatory processes on synaptic transmission remain debated (Pawlu et al., 2004; Balaji and Ryan, 2007; Alabi and Tsien, 2013).

Because Gβγ directly interacts with SNARE complexes, this opens new possibilities for the mechanistic effect of its signaling at presynaptic terminals. If Gβγ interacts with a subset of primed vesicles at the terminal it might alter P_r_ at just those sites or it might change the mode of vesicle fusion (Elhamdani et al., 2006; Alabi and Tsien, 2013). If the synapse releases just one vesicle – if it is univesicular (Maschi and Klyachko, 2020) – altering P_r_ changes the number of synaptic sites releasing transmitter for any given action potential. This will not alter transmitter concentration at each site. In contrast, variation in synaptic cleft neurotransmitter concentration might be caused by alterations in P_r_ within multivesicular release (MVR) synapses (Rudolph et al., 2015). Concentrations of transmitter at postsynaptic receptors might also be changed by altering the location of release sites within active zones with respect to the postsynaptic receptors (Tang et al., 2016). Finally, a change in synaptic cleft transmitter concentration may follow changes in vesicle fusion (Gandhi and Stevens, 2003). In the latter case, vesicle fusion begins with pore formation (Spruce et al., 1990), which expands to collapse the vesicle into the plasmalemma. The c-terminus of the SNARE complex protein, SNAP-25 is required for this expansion (Fang et al., 2015). Gβγ interacts with SNARE complexes at this region of SNAP-25 to compete with Ca^2+^-synaptotagmin interaction during evoked fusion (Zurawski et al., 2017). Thus, Gβγ-SNARE interaction opens the possibility that GPCRs regulate the synaptic vesicle fusion mode and cleft neurotransmitter concentrations.

Because colocalized GABA_B_ receptors modify Ca^2+^ entry and 5-HT_1B_ receptor effects are altered by Ca^2+^, this provides a mechanism for presynaptic integration. We have previously demonstrated that colocalized but different G_i/o_ GPCR coupled receptors target Gβγ to different effectors. At CA1-subicular synapses GABA_B_ receptors reduce Ca^2+^ entry and we now show – P_r_, whereas 5-HT_1B_ receptors modify cleft glutamate concentrations of evoked vesicle fusion events to allow modulation of postsynaptic receptors. Neither of these mechanisms alter action potential invasion of the recorded synapses as demonstrated by reliability of action potential-evoked presynaptic Ca^2+^ transients (Hamid et al., 2014). With Monte Carlo simulation of glutamate release at modeled synapses we reproduced experimental results supporting the hypothesis that GABA_B_ receptors alter P_r_ at univesicular synapses whereas 5-HT_1B_ receptor action is best explained by slowing glutamate release through a restricted fusion pore. This provides finer tuning of synaptic transmission than changes in P_r_ and creates a synaptic filter that selects for transmission of NMDA receptor-mediated events and trains of activity. We have now shown that GABA_B_ receptors, inhibit synaptic responses by changing P_r_ regardless of frequency. However, because they modify presynaptic Ca^2+^ entry which alters Gβγ-competition with synaptotagmin at SNARE complexes, GABA_B_ receptors cause meta-modulation by reshaping effects of 5-HT_1B_ receptors. Thus, presynaptic terminals show synaptic integration.

## Materials and Methods

### The preparation

Experiments were performed on hippocampal slices (300 µm) of 21-22-day-old male and female Sprague-Dawley rats anesthetized with isoflurane and decapitated in accordance with institutional guidelines. Hippocampi were isolated under semi-frozen Krebs Henseleit solution (in mM): 124 NaCl, 26 NaHCO_3_, 1.25 NaH_2_PO_4_, 3 KCl, 2 CaCl_2_, 1 MgCl_2_, 10 D-glucose, bubbled with 95% O_2_-5% CO_2_ and sliced using a Vibratome (Leica VT1200, Leica Microsystems Inc, Buffalo Grove IL) except for experiments in which high Ca^2+^ concentrations were used. In these experiments to prevent calcium carbonate precipitation, bicarbonate buffer was substituted with HEPES using a solution of the following composition in mM: 145 NaCl, 5 HEPES, 1.25 NaH_2_PO_4_, 3 KCl, 2 or 10 CaCl_2_, 1 MgCl_2_, 10 D-glucose, adjusted to pH 7.4 with NaOH and bubbled with O_2_-100% CO_2_. All recording were performed in a constant flow recording chamber in which the slice were held down with a harp. The recording chamber was superfused at approximately 2ml/min and maintained at 28°C.

### Electrophysiology

Subicular pyramidal neurons were whole-cell clamped by visual identification using an upright microscope illuminated via a fibre optic source. Recording use an Axopatch 200A or B amplifier (Axon Instruments). Patch pipettes (4-5 MΩ) contained solution (in mM): cesium (for voltage clamp) or potassium methane sulfonate (for current clamp) 146, MgCl_2_ 2, EGTA 5, HEPES 9.1, pH adjusted to 7.2 with CsOH or KOH. Series resistance was monitored by applying a 5 to 10 mV voltage step before each evoked synaptic response. If any series resistance change recorded during the experiment exceeded 20%, the cell was discarded from analysis. Focal stimuli (0.2 ms, 20 µA or less) were applied over CA1 axons using glass insulated monopolar tungsten microelectrodes (Fig 1A) (Hamid et al., 2014). Immediately after obtaining whole cell access for up to 15 minutes, a hyperpolarization and reduction in impedance could be sometimes be recorded on baclofen application. Baclofen activates postsynaptic G protein activated inwardly rectifying K^+^ (GIRK) currents to cause a change in postsynaptic impedance. This was not seen following CP93129 application. Nevertheless, in all recordings, including during those with baclofen, postsynaptic input impedance was tested with current pulses throughout the entirety of the experiment (Fig. 4). However, the baclofen evoked reduction in impedance was lost after the initial 15 min period of recording. All experimental recordings of EPSCs were made after this initial period of recording and no effect of baclofen on neuron impedance was recorded during any voltage clamp experiments. Whole cell impedance correction was not used during the experiment to avoid changes causing uncontrolled alterations in apparent EPSC amplitudes over the long recording period.

**Figure 1.**
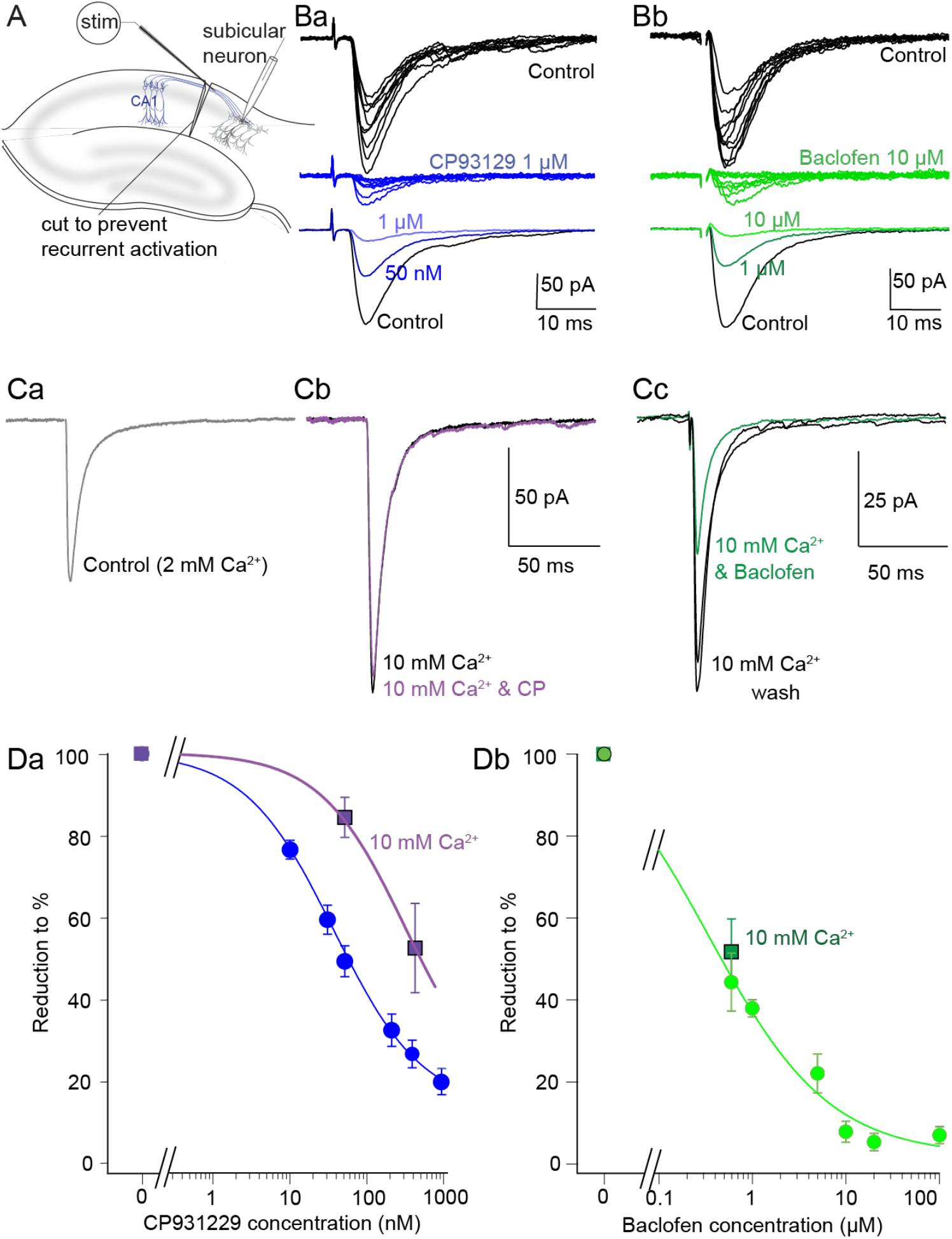
Presynaptic 5-HT_1B_ and GABA_B_ receptors inhibit evoked transmission from CA1 pyramidal neurons. A) Subicular pyramidal neurons were whole cell patch clamped. CA1 pyramidal axons were stimulated with a monopolar, glass-coated tungsten microelectrode. The tissue was cut at the CA1/subiculum boundary to prevent recurrent excitation. Ba) Ten consecutive AMPA receptor-mediated EPSCs from a cell recorded in bicuculline (5µM) and D-AP5 (50 µM) in control (black) and in CP93129 (blue). Means of responses in control (black) 50 nM (dark blue) and 1 µM CP93129 (blue) are shown below. Bb) Similar evoked EPSCs were recorded in control (black) and baclofen (10 µM) green and means of EPSCs in control and in baclofen (1µM, dark green) and 10 µM (green). C) Synaptic responses recorded in control (Ca) after addition of 10 mM CaCl_2_ (Cb, black) and with CP93129 (50 nM, purple). In raised extracellular Ca^2+^ (Cc) baclofen 1µM inhibited the synaptic response as for controls. D) Concentration responses for (Da) CP93129 in control saline (blue circles) and in 10 mM CaCl2 (purple squares) and (Db) Baclofen in control saline (light green circles) and in 10 mM CaCl_2_ (dark green square).

### Expression of iGluSnFR in hippocampus

An AAV5 containing the glutamate sensor iGluSnFr under a human synapsin promoter was intracerebrally injected in rats at postnatal days 21-22. pAAV.hSyn.iGluSnFr.WPRE.SV40 was a gift from Loren Looger (Addgene plasmid # 98929). Briefly, rats were anesthetized with isoflurane during the stereotaxic surgery and a volume of 0.2 uL of iGluSnFr was injected in the dorsal hippocampus at the following coordinates: A/P: −4.0, M/L: +/−2.7, and D/V: −1.8. Hippocampal slices were prepared from these animals 7 and 14 days after the surgery using the same procedures as described for electrophysiological experiments.

### Imaging Experiments

Line-scan confocal imaging as previously described (Hamid et al., 2019) was used for Ca^2+^ transient recording in presynaptic varicosities following single action potentials stimulated with somatic whole-cell electrodes. Alexa 594 hydrazide was excited at 568 nm. Fluo-5F was separately excited (488 nm) and imaged in bandpass (510-560 nm). Images were taken separately to ensure no cross-channel bleed-through. This was confirmed with neurons filled with only one dye. No bleed-through image was discernible for either dye to the incorrect channel. Varicosities were identified 20-35 mins after whole cell access by imaging the Alexa 594 hydrazide dye and tracking the axon from the filled soma to its projection into the subiculum. Ca^2+^ transients at these varicosities were imaged in line scanning mode (500 Hz) for up to 1 s during stimuli to the soma to evoke action potentials. Image analysis was performed within ImageJ and images are represented as linear with the applied LUT mapping.

For Lattice Light Sheet Microscopy (LLSM) hippocampal slices were placed on cover slips previously coated with 50mg/ml 300,000 MW poly-lysine in a perfused recording chamber under the LLSM objective lenses (Fig.6A). The LLSM was custom designed in our laboratory to allow simultaneous electrophysiology, but based on a Janelia Farms design (Chen et al., 2014). A twisted pair Nichrome stimulating electrode was positioned at the same location as electrophysiological experiments. Stimulation (200µs, 5 s interval) evoked fluorescence transients stochastically across the 51 x 51 µm field of view. We imaged transients using LLSM at frame rates of up to 80 Hz in single planes across multiple hotspots in a single axon. Transients were imaged in sequential LLSM frames at the same z position.

### Experimental Design and Statistical Analyses

For all values of EPSC amplitude or fits to the EPSC rise and decays, the mean was taken of at least 10 evoked traces of EPSCs for each condition for each recorded neuron. *Student* paired two-tailed t-tests were used to calculate the significance of simple paired data sets. Comparison of multiple data sets under similar conditions were made using one way ANOVA, and if necessary followed with a Post Hoc Tukey HSD test for significance of each manipulation against all others. Results are presented as numbers of recordings (*n*), the values of the means ± standard error of the mean, and for each statistical analysis are presented as degrees of freedom, F or t statistic, degrees of freedom, and absolute value of probability (*p*). Post Hoc tests following significance obtained from an ANOVA give maximum values of *p* from tables. All values of *p* were from two-tailed tests. All replicates are from different neuronal recordings in different hippocampal slices. Significance is indicated in figures over box plots and histograms as follow: * p < 0.05, ** p <0.01, *** p < 0.001.

### Monte Carlo simulations

Monte Carlo simulations were applied to neurotransmitter release through an expanding model of a synaptic vesicle fusion pore within a simulated synapse. Simulations were run in the MCell environment (Kerr et al., 2008) in which a simple 3D mesh model of a synaptic cleft was created. The cleft was modeled as disc with a 300 nm diameter and a thickness of 20 nm. Neurotransmitter release was modeled from from 1 to 3 vesicles with internal diameters of 25 nm fusing with expanding pores growing from 0.4 nm to complete opening to the diameter of the vesicle over a period varying from 200 µs to over 20 ms. An example of the sequence of frames for a single pore and its expansion and loss of glutamate to the synaptic cleft is shown in Multimedia Video 1. Glutamate escape from the simulated synaptic cleft was slowed by limiting the thickness of the cleft at its edges. This was varied systematically to result in a glutamate decay to 10% of peak in 1 ms (Clements et al., 1992; Asztely et al., 1997) following the release of 5000 glutamate molecules. After escape from the cleft, glutamate moleculas were removed from the model by simulating pumps external to the synaptic cleft. The vesicles were modeled containing from 700 to 20000 glutamate molecules. Kinetic models of glutamate receptors were placed in the postsynaptic disc (20 NMDA and 20 AMPA receptors) enabling simulation of the binding, activation, desensitation, inactivation and unbinding of these receptors. Kinetic models of the receptors are shown in Figure 7. Model parameters including those of the receptor kinetic models used are included in table 1. Animations of these simulations are shown in Multimedia Video 1 and 2).

**Table 1.** Monte Carlo Simulation Parameters and Sources. Table shows parameter values and sources for the monte carlo simulations in figures 7 and 8

| Parameter | value | source |
| --- | --- | --- |
| Glutamate diffusion constant (cm<sup>2</sup>.s<sup>-1</sup>) | 3 x 10<sup>-6</sup> | (Rusakov and Kullmann, 1998) |
| AMPA receptor parameters (glutamate) |  |  |
| k1 (M<sup>-1</sup>.s<sup>-1</sup>), k-1 (s<sup>-1</sup>)<br>k-2 (s<sup>-1</sup>)<br>γ<sub>0</sub>, δ<sub>0</sub> (s<sup>-1</sup>)<br>γ<sub>1</sub>, δ<sub>1</sub> (s<sup>-1</sup>)<br>γ<sub>2</sub>, δ<sub>2</sub> (s<sup>-1</sup>)<br>β, α (s<sup>-1</sup>) | 2 x 10<sup>7</sup>, 9 x 10<sup>3</sup><br>0.41<br>1, 3 x 10<sup>-3</sup><br>7.6, 1.8 x 10<sup>3</sup><br>35, 200<br>8 x 10<sup>3</sup>, 3.1 x 10<sup>3</sup> | (Robert and Howe, 2003) |
| (kynurenate) |  |  |
| k3 (M<sup>-1</sup>.s<sup>-1</sup>)<br>k-3 (s<sup>-1</sup>) | 5 x 10<sup>7</sup><br>9000 | (Diamond and Jahr, 1997) |
| NMDA receptor parameters |  |  |
| on, off<br>d1+, d1-<br>d2+, d2-<br>f+, f-<br>s+, s- | 95 x 10<sup>5</sup>, 29<br>0.5, 45<br>70, 2.8<br>1557, 182<br>89, 135 | (Banke and Traynelis, 2003) |
| Mesh properties |  |  |
| Internal vesicle diameter (nm)<br>Pore length (nm)<br>Minumum pore diameter (nm)<br>Synaptic cleft thickness (nm) | 25<br>8<br>0.4<br>20 | (Hu et al., 2008)<br>(Vardjan et al., 2007)<br>(Savtchenko and Rusakov, 2007) |
| Synaptic cleft diameter (nm)<br>Thickness of cleft edge diffusion barrier (nm) | 300<br>1.7 | (Nimchinsky et al., 2002)<br>(Rusakov and Kullmann, 1998) |

### Code Accessibility

Simulation parameter files are available at: https://alford.lab.uic.edu/GPCRs.html and at ModelDB for requested access at http://modeldb.yale.edu/266839 prior to publication of this work. This will be made public on publication.

The authors declare that experimental data supporting the findings of this study are available within the paper.

## Results

### GABA_B_ and 5-HT_1B_ receptors inhibit synaptic transmission from CA1 to subiculum

5-HT_1B_ and GABA_B_ receptors have previously been shown to be co-expressed in CA1 axon presynaptic terminals. (Boeijinga and Boddeke, 1996; Bonaventure et al., 1998; Lein et al., 2007; Hamid et al., 2014) where both receptors inhibit synaptic transmission, but by different mechanisms. We have previously shown that GABA_B_ receptors inhibit Ca^2+^ entry, while 5-HT_1B_ receptors cause Gβγ to act directly at the SNARE complex (Hamid et al., 2014). The amount of inhibition of Ca^2+^ entry by GABA_B_ receptors in that study was sufficient to account for the effect of the GABA_B_ receptor agonist, baclofen on synaptic transmission. In contrast, the effect of the 5-HT_1B_ receptor agonist, CP93129 was eliminated by treatment of the tissue with Botulinum A toxin to remove the c-terminal 9 residues from SNAP-25 and in a further study (Zurawski et al., 2019) genetic manipulation of the c-terminal of SNAP-25 also prevented 5-HT_1B_ receptor modulation of neurotransmitter release. We also showed in those studies no loss of transmission between soma and axons. This was indicated by Ca^2+^ transients, that were unnaffected by 5-HT_1B_ receptors and reduced in amplitude, but not prevented by GABA_B_ receptors (Hamid et al., 2014).

To investigate the effects of GABA_B_ and 5-HT_1B_ receptors on neurotransmitter release, we whole cell voltage clamped subicular pyramidal neurons (Fig. 1A) and stimulated CA1 axons with glass coated tungsten microelectrodes at low intensity to ensure focal stimulation (5-20 µA, 200 µs). Recorded events were of low amplitude (Fig 1B): approximately 10 times the unitary amplitude determined from spontaneous release in TTX shown in earlier work (Hamid et al., 2014), which ensured that responses were monosynaptic. To prevent polysynaptic firing by recurrent excitation of CA1 neurons, the tissue was cut at the CA1-subicular boundary. We recorded responses in bicuculline (5µM) and 2-amino-5-phosphonopentanoic acid (AP5, 50 µM) to isolate AMPA receptor-mediated currents. We applied the selective 5-HT_1B_ receptor agonist CP93129. Effects of a saturating concentration (1 µM) and at a concentration approximately the EC50 (50 nM) are shown (Fig. 1Ba, Fig. 1Da for the concentration response). CP93129 reduced the EPSC peak amplitude to 19.6 ± 3.9% at a saturating dose (1µM, n=8; EC50=0.033 ± 0.04µM). These EPSCs are also inhibited by the GABA_B_ receptor agonist, baclofen to 7.2 ± 2.3% of control (100µM baclofen, n=5, EC50=0.35 ± 0.18µM) (Fig. 1Bb, Db).

We have previously shown that 5-HT_1B_ receptors target Gβγ to SNARE complexes in CA1 terminals (Hamid et al., 2014), and Gβγ competes with Ca^2+^-synaptotagmin binding to SNARE complexes. In lamprey this may confer Ca^2+^ dependency on inhibition (Yoon et al., 2007). To test for this in CA1 subicular synapses, we recorded EPSCs in HEPES buffered Ringer (to prevent Ca^2+^ precipitation). Increasing CaCl_2_ concentrations (10mM) enhanced the EPSC (Fig. 1Ca,b), but CP93129 now caused significantly less inhibition (Fig. 1Cb, Da; at 50nM, paired Students t test, t(15)=6.00, p=0.00001 n=13 and 8; and at 400nM t(4)=2.46, p=0.03, n=10 and 4). The EC50 of CP93129 increased from 33 ± 4 nM to 320 ± 19 nM. In contrast, baclofen (600nM) inhibition was unaffected by high Ca^2+^ concentration (t(16)=0.88, p=0.38; Fig. 1Cc, Db). This latter result implies that 10 mM Ca^2+^ does not cause saturating Ca^2+^ entry with respect to neurotransmitter release, allowing reduced Ca^2+^ entry by baclofen to reduce P_r_.

### Paired pulse ratios are modified by GABA_B_ receptors but not by 5-HT_1B_

Colocalized GABA_B_ and 5-HT_1B_ receptors act at different targets – Ca^2+^ channels or the SNARE complex respectively – therefore their activation may evoke different effects on vesicle fusion. We measured paired pulse ratios (PPRs) to determine whether these receptors cause inhibition by a change in P_r_ (Dobrunz and Stevens, 1997). As a control we raised the extracellular Mg^2+^ concentration (Del Castillo and Katz, 1954) from 1 to 4mM to reduce presynaptic Ca^2+^ entry and therefore P_r_. In 4mM Mg^2+^, the mean amplitude of the first EPSC was reduced (to 42 ± 3% of control n=14, Fig. 2A), and the PPR (EPSC2/EPSC1) was increased by a factor of 1.64 ± 0.09, n=14 (quantified in Fig. 2E, red) demonstrating a Ca^2+^ dependent reduction in P_r_.

**Figure 2.**
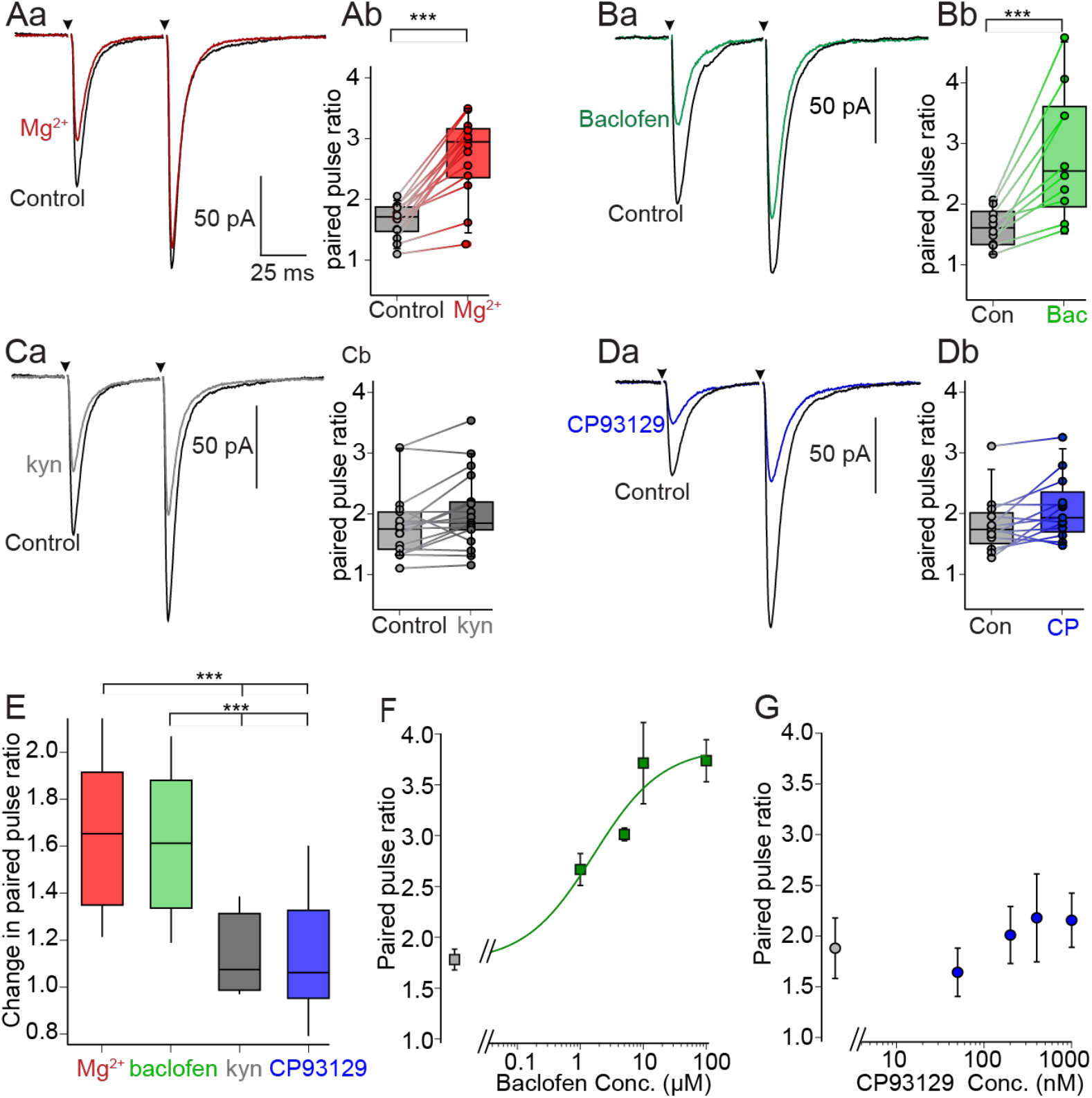
P_r_ is unaffected by 5-HT_1B_ but reduced by GABA_B_ receptor activation. A) Paired pulse stimuli applied to CA1 axons (50ms intervals). (Aa) Means of 10 EPSCs. Extracellular [Mg^2+^] was raised from 1 to 4mM to reduce Ca^2+^ entry (red) and thus P_r_. This reduced the mean amplitude of the first EPSC more than the second. The effect of high Mg^2+^ on the paired pulse ratio (PPR) is quantified to show scatter plots of PPRs in control and high Mg^2+^ for each cell and box plots of the same data (Ab). B) A similar experiment but with addition of the GABA_B_ receptor agonist, baclofen. (Ba, 600 nM), mean EPSCs show an increase in PPR quantified in Bb as for Ab. C) A negative control using a glutamate receptor antagonist, kynurenate (200μM; grey). This gave no change in PPR (Ca, quantified in Cb). D) Similar experiment but with the 5-HT_1B_ receptor agonist, CP93129 (50nM; blue). This again gave no change in PPR (Da, quantified in Db). E) Changes in PPRs normalized to pre-treatment controls shown as box plots (1.0 = no change, boxes 25 and 75% with means, whiskers to 10 and 90%) for all recordings in the conditions shown in (A-D). F) The effect of baclofen concentration vs. normalized PPR was plotted. This is concentration dependent. G) The effect of CP93129 concentration on PPR was plotted. This was not concentration-dependent

We inhibited EPSCs with baclofen (600nM, ∼ half maximal concentration; Fig. 2B) which reduced the first EPSC to 40 ± 6% and enhanced PPRs (by 1.62 ± 0.1, n=10; quantified in Fig. 2E, green) demonstrating an effect of GABA_B_ receptors on P_r_, consistent with our earlier work showing their effect on presynaptic Ca^2+^ entry (Hamid et al., 2014). The effect on PPR was concentration dependent (Fig. 2F) with a similar concentration response to the effect on EPSC amplitude (Fig 1Db).

As a negative control, AMPA receptor antagonism with half-maximal concentrations of the glutamate receptor antagonist kynurenate (200µM) showed no effect on PPRs. (Fig. 2C. The EPSC2/EPSC1 ratio was 1.13 ± 0.04 (n=19) of control (quantified Fig. 2E, gray). This is expected for postsynaptic targets. CP93129, inhibits CA1-subicular EPSCs presynaptically (Zurawski et al., 2019). Nevertheless, we now show a half-maximal CP93129 (50nM) concentration, which reduced the first EPSC to 54 ± 4% of control, did not alter PPRs (PPR in CP93129 was 1.13 ± 0.07 of control; Fig. 2D,E, blue; n=14). This lack of effect of CP93129 on PPRs was consistent over the full concentration response range of CP93129 on EPSCs (Fig. 2G).

A one way ANOVA to test effects of raised Mg^2+^ concentration, of baclofen, kynurenate and of CP93129 on PPRs (data from Fig. 2E), showed significance, ANOVA single factor F(3,53) = 16.85, p=8 x 10^−8^). Post-hoc Tukey HSD analysis revealed no significant difference between effects of raised Mg^2+^ and baclofen (p>0.1) and none between effects of kynurenate and CP93129 (p>0.1). However, the effect of CP93129 was significantly different than lowering P_r_ with raised Mg^2+^ (p<0.001), or baclofen (p<0.001). Thus, effects of 5-HT_1B_ receptors are paradoxical, they are presynaptic but do not alter P_r_, whereas GABA_B_ receptors which are also presynaptic do alter P_r_.

### 5-HT1B but not GABA_B_ receptors enhance low affinity block of AMPA EPSCs

We investigated whether presynaptic receptors alter synaptic cleft glutamate concentrations at postsynaptic AMPA receptors. Partial block of EPSCs by a low affinity antagonist is caused by antagonist binding and unbinding. If the antagonist unbinds within the time-course of glutamate in the synapse, antagonist bound receptors will then become available for either glutamate or antagonist binding. Therefore, fractional low affinity antagonism of the AMPA receptor EPSC will depend on the glutamate concentration (Clements et al., 1992). EPSCs evoked by low glutamate cleft concentrations will be more completely antagonized than those at high concentrations.

To first determine any effect of a reduction in P_r_ on cleft glutamate concentration, we recorded AMPA receptor EPSCs. The effect of lowering P_r_ with high Mg^2+^ was tested against the efficacy of low affinity kynurenate antagonism. We compared kynurenate efficacy in control (1mM) and high (4mM) Mg^2+^ concentrations in the same cell and stimulus intensity, because it is not clear that the mean control cleft glutamate concentrations were identical from cell to cell. After obtaining baseline EPSCs, we superfused the slice with kynurenate (200μM) to obtain a partial block of the EPSC (in the example to 70% of control; Fig. 3Aa, gray). Kynurenate was removed to re-measure control EPSCs. The Mg^2+^ concentration was then increased to 4mM to reduce P_r_ (EPSC reduced to 69% of control, Fig. 3Aa, red). Kynurenate (200µM) was reapplied in 4mM Mg^2+^ to measure its efficacy at a lower P_r_. The EPSC in 4 mM Mg^2+^ was reduced by the same proportion as in control (to 67% of response in 4mM Mg^2+^, Fig. 3Aa, purple). To compare inhibition by kynurenate in control and 4mM Mg^2+^ the responses in 4mM Mg^2+^ alone and combined with kynurenate were scaled to control amplitudes (Fig. 3Ab) showing similar proportionate inhibition. In repeat experiments the drug sequence was randomized. For all cells, in 1mM Mg^2+^, kynurenate (200µM) reduced EPSCs to 44 ± 4% of control, and in 4mM Mg^2+^ kynurenate similarly reduced the EPSC to a not significantly different 43 ± 4% (n=10; Fig. 3Ac, purple; paired Students t test, t(9)= 0.39, p=0.7).

**Figure 3.**
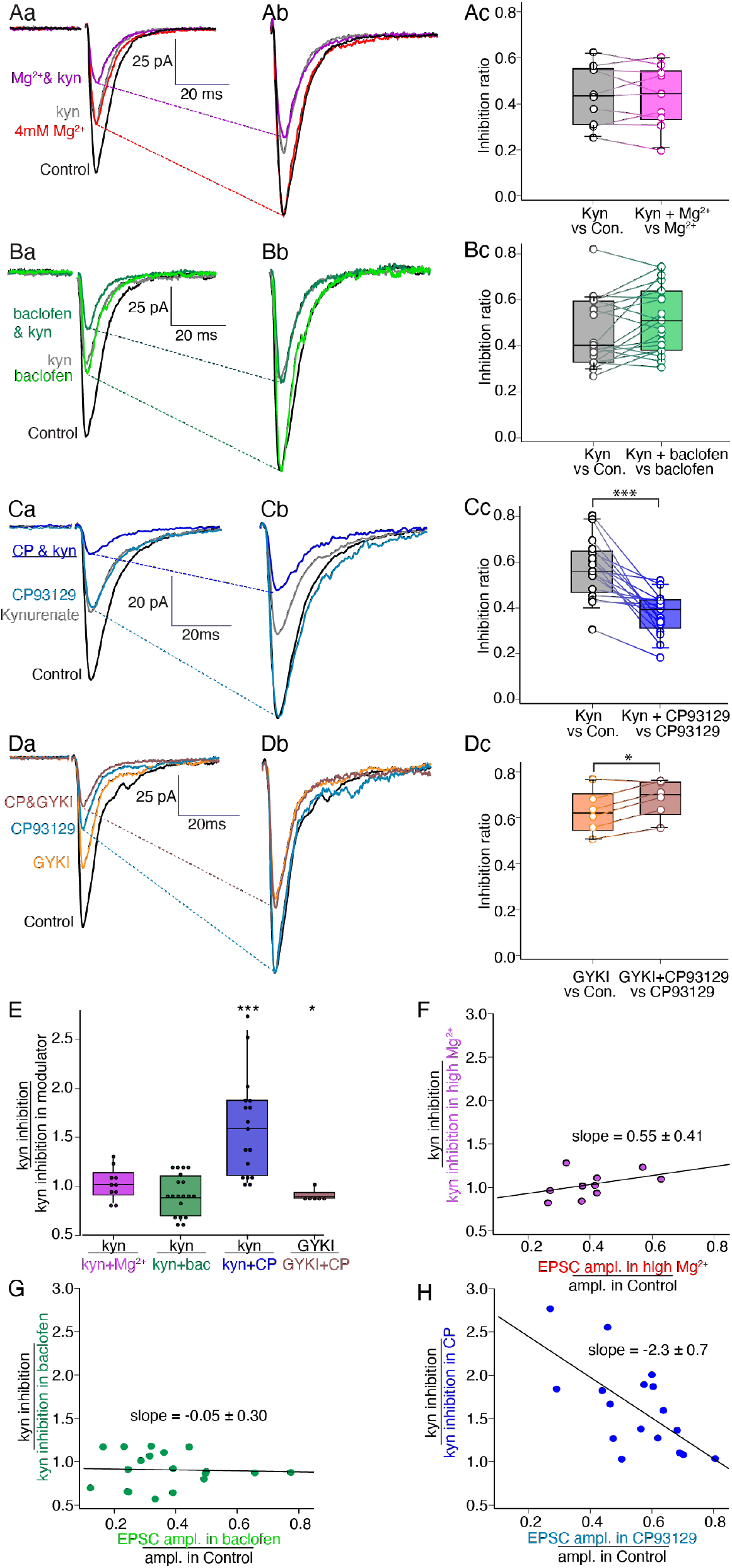
5-HT_1B_ but not GABA_B_ receptors increase the potency of a low affinity AMPA receptor antagonist-mediated block of EPSCs. A) Effect of raised Mg^2+^ on EPSC inhibition by a low affinity competitive antagonist, kynurenate. (Aa) Kynurenate (200μM; gray) partially inhibited EPSCs, shown as means of 10 traces. After kynurenate wash, 4 mM Mg^2+^ (red), to decrease P_r_ reduced EPSCs. Kynurenate 200μM was then combined with high [Mg^2+^] (purple). (Ab) For clarity, traces in 4mM Mg^2+^ scaled to control. (Ac) Kynurenate inhibited EPSCs to a similar ratio in control and 4mm Mg^2+^. Inhibition ratios plotted and overlaid on boxplots (boxes 25 and 75% with means, whiskers to 10 and 90%) Ba) Kynurenate (200μM; gray) reduced EPSCs. After kynurenate wash, baclofen (600 nM; light green) reduced the EPSC. Kynurenate (200 µM) with baclofen (600 nM; dark green) reduced the EPSC similarly to baclofen alone. (Bb) Reponses in baclofen scaled as for Ab. (Bc) data from all cells. Ca) kynurenate (200μM; gray) reduced EPSCs. After wash, CP93129 (50nM; light blue) reduced the EPSC. Kynurenate (200 µM) with 50nM CP93129 (blue) reduced the EPSC to a greater proportion than CP93129 alone. (Cb) Reponses in CP93129 scaled as for Ab. (Cc) data from all cells. D) Non-competitive AMPA receptor antagonist GYKI used as control. (Da) GYKI (10μM; orange) reduced EPSCs. After wash, CP93129 (50nM) inhibited EPSCs (blue). Co-application of 10μM GYKI with 50nM CP93129 (pink) reduced EPSCs as controls. (Db) Responses in CP93129 scaled as above. (Dc) data from all cells. E) Box plots overlaid with individual cell data comparing ratio of inhibition by kynurenate or GYKI alone, to inhibition in kynurenate plus high Mg^2+^ (pink), baclofen (green) or CP93129 (blue) or to GYKI and CP93129 (brown). A value of 1 indicates the modulatory agonist had no effect on kynurenate or GYKI potency, higher values show enhanced potency. F) Change in kynurenate inhibition caused by 4mM Mg^2+^ (data from A) plotted against initial efficacy of 4mM Mg^2+^ for each neuron. The slope was indistinguishable from zero. G) Similar effect by baclofen on kynurenate inhibition (B) plotted against initial efficacy of baclofen in each neuron. Slope was again indistinguishable from zero. H) Change in kynurenate inhibition by CP93129 (C) plotted against the initial efficacy of CP93129 in each neuron. Slope (−2.3) showed correlation (R^2^ =0.4).

We then assayed the efficacy of kynurenate inhibition before and in baclofen which will also alter P_r_. In the example, kynurenate (200μM) reduced the mean EPSC amplitude to 62% of control (Fig. 3Ba, gray). In baclofen (600 nM; Fig. 3Ba, green) the inhibited EPSC was re-probed with kynurenate (200µM), which now reduced the EPSC to 61%% of its amplitude in baclofen alone (Fig. 3Ba, dark green). Responses were again scaled to compare the effect of kynurenate in controls and in baclofen (Fig. 3Bb). In 19 experiments, kynurenate inhibited control responses to 46 ± 3% and the inhibitory effect was slightly but signficantly less in baclofen, inhibiting to 51 ± 3 % (Fig. 3Bc; green; paired Students t test, t(18) = 2.19, p = 0.04).

We similarly assayed the efficacy of kynurenate block before and in CP93129, which did not alter P_r_. In the example, kynurenate (200μM) reduced peak the EPSC amplitude to 55% of control (Fig. 3Ca, gray). After removal of kynurenate, EPSC amplitudes were recorded to reconfirm control responses. CP93129 (50nM) was applied (Fig. 3Ca, light blue), and the CP93129 inhibited EPSC was re-probed with kynurenate (200µM), which now reduced the EPSC to 38% of its amplitude in CP93129 alone (Fig. 3Ca, blue). Responses were again scaled to directly compare the effect of kynurenate in controls and in CP93129 showing an enhancement in inhibition (Fig 3Cb). In 17 experiments, kynurenate inhibited control responses to 57 ± 3% and responses in CP93129 to 37 ± 2%. This latter inhibitory effect was signficantly greater than the effect of kynurenate alone (Fig. 3Cc; (paired Students t test, t(16) = 5.56, p = 2.1×10^−5^).

As a negative control for the effect of CP93129 in kynurenate, we replaced the competitive antagonist (kynurenate) with a non-competitive AMPA receptor antagonist, GYKI52466 (GYKI). GYKI should inhibit EPSCs similarly, regardless of glutamate concentration because it does not compete with glutamate (Fig 3D). Mean inhibition by GYKI in control was to 63 ± 4% and the efficacy of GYKI was slightly but significantly reduced in CP93129 (n=6, to 68 ± 4% of response in CP93129, paired Students t test, t(5)=3.85, p=0.011). This result is small but may imply a slight bias in analysis recordings of EPSC amplitudes as their amplitude is reduced to very small values by combined ligands. Recording noise may contribute slightly to the computed result because the EPSC measurement algorithm searched for a smoothed positive maximum. Thus, in very small or results with a failure there may be a small positive bias in the result. The result may indicate that we have slightly underestimated the effect of 5-HT_1B_ receptors on cleft glutamate concentrations, but does will not change the conclusion. This bias may also underlie the slight but not significant relative reduction in effect of kynurenate seen in baclofen (Fig 3 Bc).

The effects of high Mg^2+^, baclofen and CP93129 on the efficacy of kynurenate and the effect of CP93129 on the efficacy of GYKI on the inhibition of the synaptic responses were compared to one another by comparing the ratios of effects of antagonist (kynurenate or GYKI) with the same antagonist combined with either 4 mM Mg^2+^, with baclofen, or with CP93129 (Fig. 3E). All four experiments were compared with a one way ANOVA showing substantial significance (F(3,48) = 15.88, p = 1.4 x 10^−6^). Post hoc Tukey’s HSD analysis revealed that the the effect of baclofen was not significantly different than the effect of high Mg^2+^ on the efficacy of kynurenate inhibition (p > 0.1), This is consistent with inhibition mediated by a change in P_r_ of univesicular events at each synapse. In contrast, the effect of CP93129 on kynurenate-mediated inhibition was significantly greater than that of Mg^2+^ (p < 0.005), baclofen (p < 0.001) and of the effect of CP93129 on GYKI inhibition (p < 0.005).

Single raised concentrations of Mg^2+^ (4mM) or single concentrations of baclofen (600nM), or CP93129 (50nM) show varying degrees of inhibition from cell to cell. For example, initial effects of 50 nM CP93129 reduced the EPSC from as little as to 0.8 of control to as much as 0.3. If this variance reflects the efficacy of the agonist on cleft glutamate concentration it will correlate with amplification of agonist inhibition by kynurenate (shown in Fig. 3E). To test this we compared the efficacy of the agonist alone in each neuron (agonist/control), to the change in agonist efficacy by kynurenate [(kynurenate/control)/(agonist/kynurenate+agonist)]. We first plotted these for effects of high Mg^2+^ (Fig. 3F). There was no correlation between inibition by 4mM Mg^2+^ with the subsequent effect of Mg^2+^ on kynurenate inhibition. This is consistent with a Mg^2+^-induced change in P_r_ at univesicular synapses causing no change in cleft glutamate concentration regardless of the initial efficacy of raised Mg^2+^. Similarly, there is no correlation between the efficacy of baclofen and its effect on subsequent kynurenate block (Fig. 3G). However, there is a strong correlation between the CP93129 efficacy and its amplification of kynurenate inhibition (Fig 3H), indicating that the greater the effect of presynaptic 5-HT_1B_ receptors, the lower it drives cleft glutamate concentration and therefore the larger inhibitory effect of kynurenate. We conclude that at CA1 presynaptic terminals synapses changes in P_r_ do not alter cleft glutamate concentrations, but 5-HT_1B_ receptors do.

It was important to determine that synaptic responses were not impacted by changes in whole cell recording access during the experiments. In all experiments, we monitored whole cell access by applying a small (5 mV, 20 ms) test pulse before each evoked EPSC was recorded. An example is shown of this recording throughout a sequential application of kynurenate, and CP93129 and then the drugs combined simultaneously measuring effects on synaptic responses (Fig. 4A,B), while simultaneously monitoring whole cell series resistance calculated from double exponentials fitted to the settling of the step current (Fig. 4C). Experiments were terminated and data not used if access resistance changed by more than 20%. For the examples used for analysis (Fig 4D), no signficant difference was seen for access impedance (ANOVA single factor F(3, 64) = 0.065, p = 0.97) or holding current (F(3, 64) = 0.18, p = 0.91) when comparing between periods of ligand applications.

**Figure 4.**
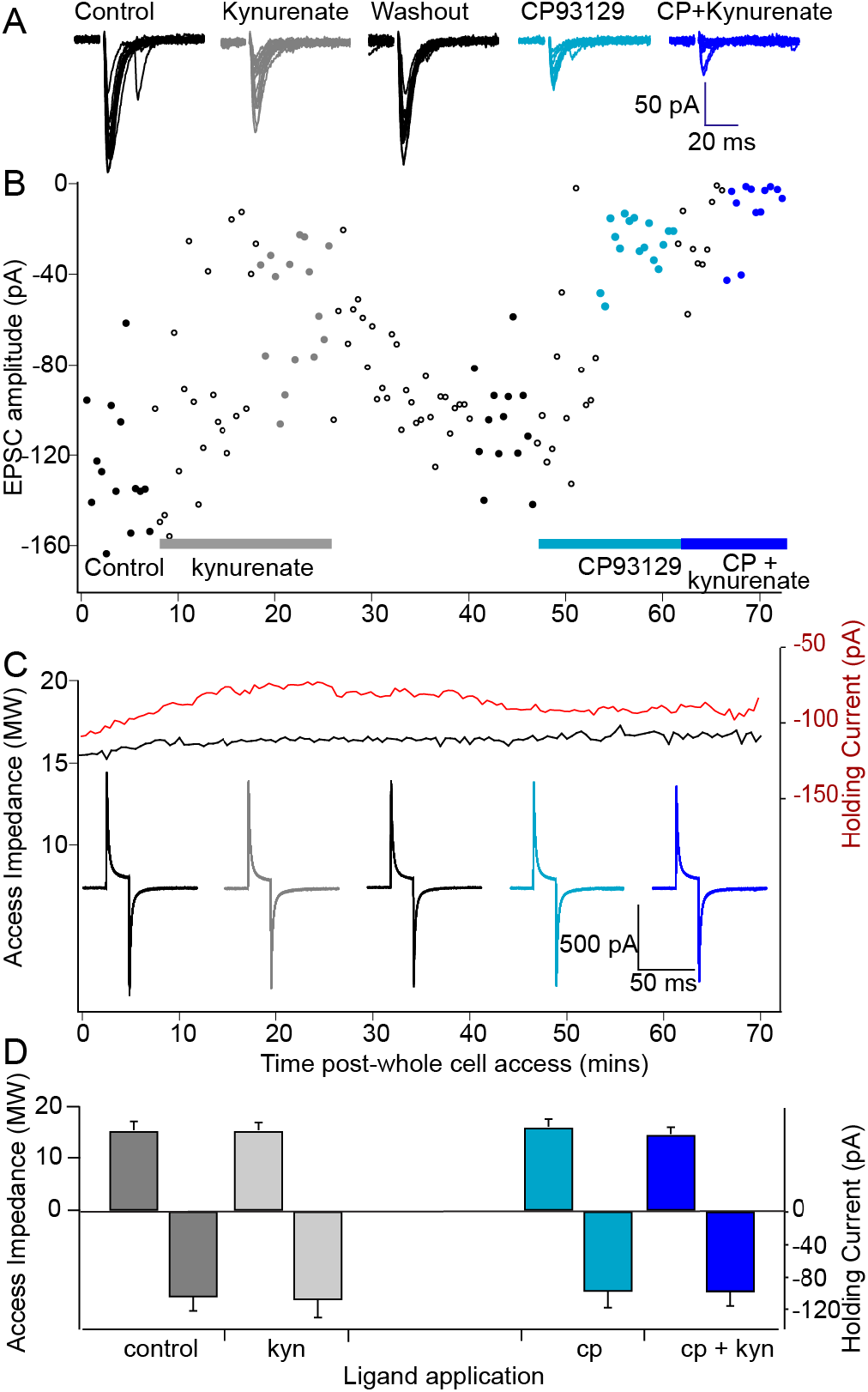
Stability of Recording Access. A) Sequential recordings of evoked AMPA receptor-mediated EPSCs during recording through each drug condition to show recording stability. Individual EPSCs are shown for one recording in which the effects of CP93129 and its amplification by kynurenate were tested. B) Graph of peak amplitudes of all EPSCs were recorded. Closed circles are measured responses corresponding to the EPSCs shown in (A). Open circles during washin in or washout of the drugs were not measured. Colors correspond to EPSCs in (A). C) Graphs show whole cell access impedance (black), and holding current (red) throughout the experiment. Whole cell access impedances (*R_a_*) were calculated from double exponentials fit to the decays from small depolarizing current steps applied before each EPSC was stimulated (Sigworth, 1983). Example responses to voltage steps at each drug application are shown. Color as as for (A). D) From all cells, histograms show means ± sem for access impedance (positive bars) and holding currents (negative bars) during control, application of kynurenate, of CP93129 and kynurenate+CP93129. Colors during ligand applications are the same throughout the figure.

### 5-HT1B but not GABA_B_ receptors differentially inhibit AMPA and NMDA EPSCs

Glutamate activates receptors with varying sensitivities: for example, AMPA, NMDA and metabotropic glutamate receptors have different affinities for glutamate and different numbers of glutamate binding sites. Consequently, changes in cleft glutamate concentration may differentially alter their activation (Choi et al., 2003; Schwartz et al., 2007; Gerachshenko et al., 2009). We compared 5-HT_1B_ and GABA_B_ receptor inhibition on AMPA and NMDA receptor-mediated synaptic responses. AMPA receptor-mediated EPSCs were recorded as before. We recorded NMDA receptor-mediated synaptic responses in bicucullline (5µM), and 2,3-dioxo-6-nitro-7-sulfamoyl-benzo[f]quinoxaline (NBQX; 5µM) at a holding potential of −35 mV. Baclofen (600nM) equally inhibited AMPA and NMDA components of EPSCs (Fig. 5A; to 49 ± 6 %, n=8, and to 54 ± 4 %, n=6 respectively; t(12)=0.80, p=0.44). CP93129 (50nM) inhibited AMPA EPSCs significantly more than NMDA EPSCs (Fig. 5B; to 54 ± 6 %, n=10, and to 93 ± 4 %, n=6 respectively; t(14)=7.15, p=4.9 x 10^−6^). Concentration response curves of CP93129 inhibition of AMPA and NMDA EPSCs diverged (Fig. 5C). AMPA receptor-mediated EPSCs were maximally inhibited to 20 ± 3 % of control whereas NMDA receptor-mediated EPSCs were significantly less inhibited to 75 ± 6 % (paired Students t test, t(7)=6.48, p=0.00017). We obtained similar results when we recorded NMDA and AMPA EPSCs simultaneously from the same synapses. AMPA and NMDA receptor-mediated responses were recorded simultaneously by holding the membrane potential at positive values and recording events early in the response or later. We measured AMPA receptor EPSC amplitudes within 2 ms of stimulation, whereas NMDA responses were measured during the decay after 100 ms (Fig. 5D) because these receptors show much slower decay kinetics than AMPA receptor mediated responses (Lester et al., 1990). In control conditions with no drugs, NMDA receptor EPSC amplitudes co-vary with those of AMPA receptors (Fig. 5E) indicating that the receptors are activated on the same synapses. In 6 neurons, we plotted the mean amplitudes of AMPA and NMDA receptor mediated EPSCs with increasing doses of CP93129. AMPA receptor EPSCs were inhibited to a significantly greater extent than were NMDA receptor EPSCs at all applied concentrations of CP93129 (Fig. 5F). For the approximate half maximal concentration (50 nM) t(5)=5.61, p=0.002) and for the maximal concentration (1000 nM) t(5)=7.52, p=0.0007).

**Figure 5.**
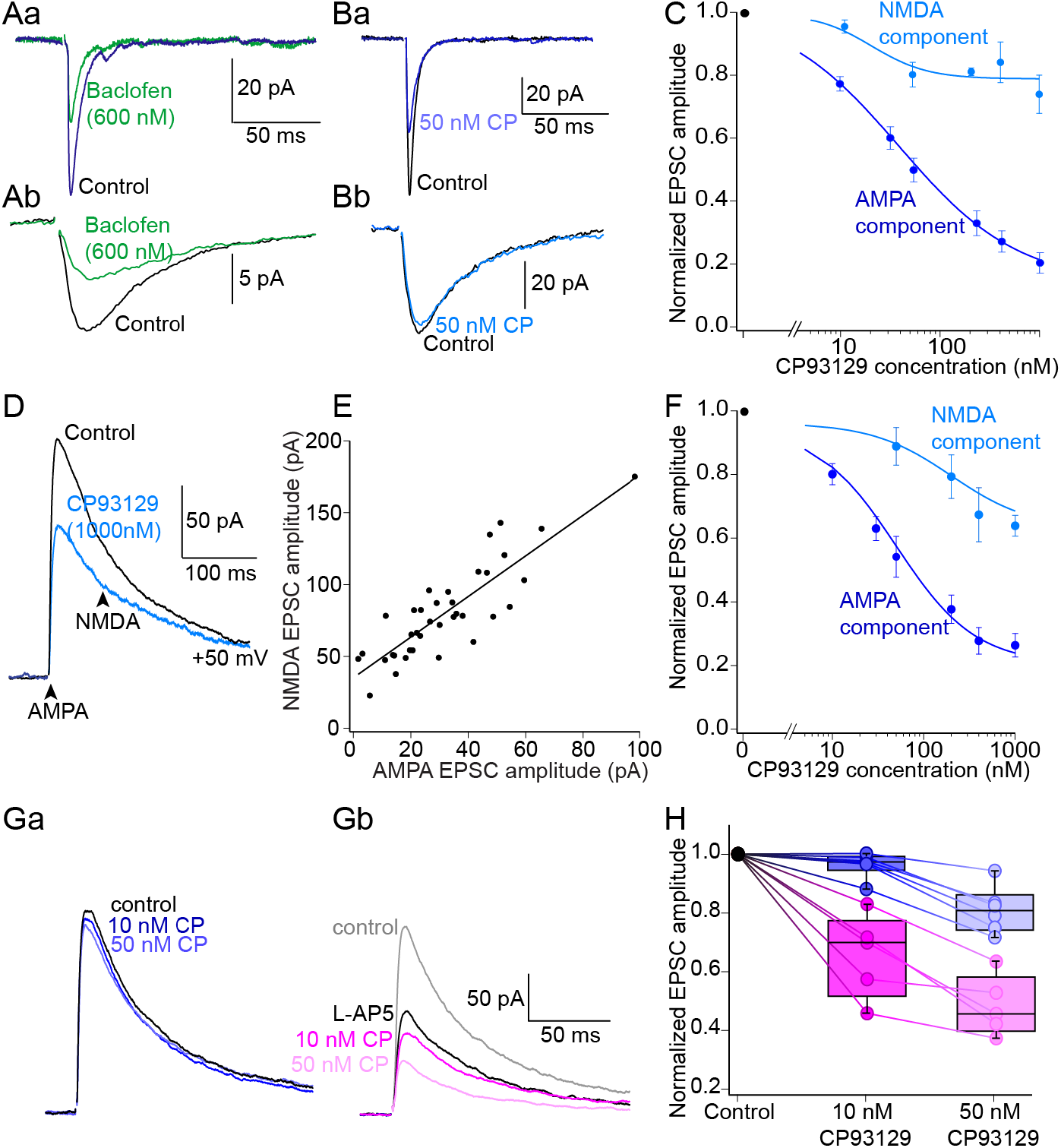
Differential inhibition of AMPA and NMDA receptor mediated EPSCs. A) In separate recordings AMPA mediated EPSCs were recorded in D-AP5 (Aa) and NMDA-mediated EPSCs in NBQX (Ab). Baclofen (600 nM; green) similarly inhibited both of these responses. B) Similar recordings were made but CP93129 (Ba, 50 nM; blue) inhibited the AMPA receptor mediated EPSC, but not the NMDA receptor mediated EPSC (Bb, light blue). C) Concentration response effects of CP93129 vs the AMPA (blue) and NMDA receptor EPSCs (light blue) show a large difference in efficacy. D) The differential effect of CP93129 on AMPA and NMDA receptor responses are found at the same synapses. Compound EPSCs without glutamate receptor antagonists recorded at a holding potential of +50mV. Early responses (2 ms post stimulus) are dominated by AMPA receptors (bottom arrow), late responses (100 ms post stimulus) by NMDA receptors. CP93129 at a saturating dose (1 µM) was applied (blue). E) For individual recordings, sequential response amplitudes of AMPA and NMDA responses were plotted against each other. These responses covaried indicating they colocalise to the same synapses. F) Mean amplitudes of AMPA and NMDA receptor mediated EPSCs during the same stimuli plotted with increasing concentrations of CP93129. AMPA receptor EPSCs were inhibited to a greater extent than were NMDA receptor EPSCs. G) CP93129 mediated inhibition of the NMDA component is amplified by the low affinity competitive antagonist (L-AP5). In control experiments (Ga) 10 and 50 nM CP93129 (blue) had little effect on isolated NMDA mediated EPSC (recorded at +50 mV in NBQX and bicuculline). L-AP5 (250 µM) partially inhibited this response (Gb, black). This inhibited response was now substantially inhibited by 10 and 50 nM CP94129 (red and pink). H) Quantitation of the data from (G) showing inhibition by CP93129 in control (blue, normalized to precontrol recordings) and after treatment with L-AP5 (pink, normalized to recordings in L-AP5 alone.

If the lack of effect of CP93129 on NMDA EPSCs is because it causes reduced cleft glutamate concentrations then an effect of CP93129 would be revealed by low affinity NMDA receptor antagonists that will amplify effects of changing glutamate concentrations at NMDA receptors (Choi et al., 2000). We recorded evoked pharmacologically isolated NMDA receptor EPSCs (n=6) at +50mV in bicuculline (5µM) and NBQX (5µM). Low doses of CP93129 (10 and 50nM) did not significantly reduce the EPSC in 6 neurons (ANOVA single factor F(2,12)=3.80, p=0.052; Fig. 5Ga, H). In a further 6 neurons the same experiment was performed in the low affinity competitive NMDA receptor antagonist L-5-amino pentanoic acid (L-AP5, 250 µM). The EPSC was reduced to 55 ± 4 % of control by L-AP5 (Fig. 5Gb, H). In L-AP5, CP93129 at both 10 and 50nM CP93129 now inhibited these partially blocked EPSCs to 66 ± 7 and 45 ± 4 % of the response in L-AP5 alone. These reductions were significant (ANOVA single factor F(2,12)=7.22, p=0.009). Thus, NMDA receptors are exposed to a similar 5-HT_1B_ receptor-mediated reduction in glutamate concentration to that seen by AMPA receptors.

### The effect of 5-HT_1B_ receptors on glutamate release

It is possible to image glutamate release using genetically engineered glutamate sensors. To achieve this we virally infected subicular neurons with iGluSnFR using an AAV1 containing iGluSnFR.WPRE.SV40 under a human synapsin promotor (pAAV.hSyn.iGluSnFr.WPRE.SV40 (AAV5) (Marvin et al., 2013) by injecting the viral vector into hippocampi of 22 day rats. In hippocampal slices from these animals epifluorescence imaging (excitation 470 nm, emission, 520 nm) revealed fluorescent subicular pyramidal neurons. The slices were placed in a custom designed imaging chamber and a stimulation electrode placed as for electrophysiogical recording. The subicular region was then imaged using LLSM (Fig 6A). The light sheet penetrated to aproximately 50 µm in the slice and time series images of single planes (512 x 512 pixels; 52 x 52 µm) were captured with a single exposure time of 10 ms and a frame rate of 80 Hz. Stimuli were applied to CA1 axons at 5 second intervals to evoke events that appeared stochastically located, varying from stimulus to stimulus across the field of view (Fig 6B). Repeated stimuli revealed multiple locations of release that were resolved over a sequence of up to 21 individual stimuli. Events were well-resolved using LLSM imaging (Fig 6C,D).

**Figure 6.**
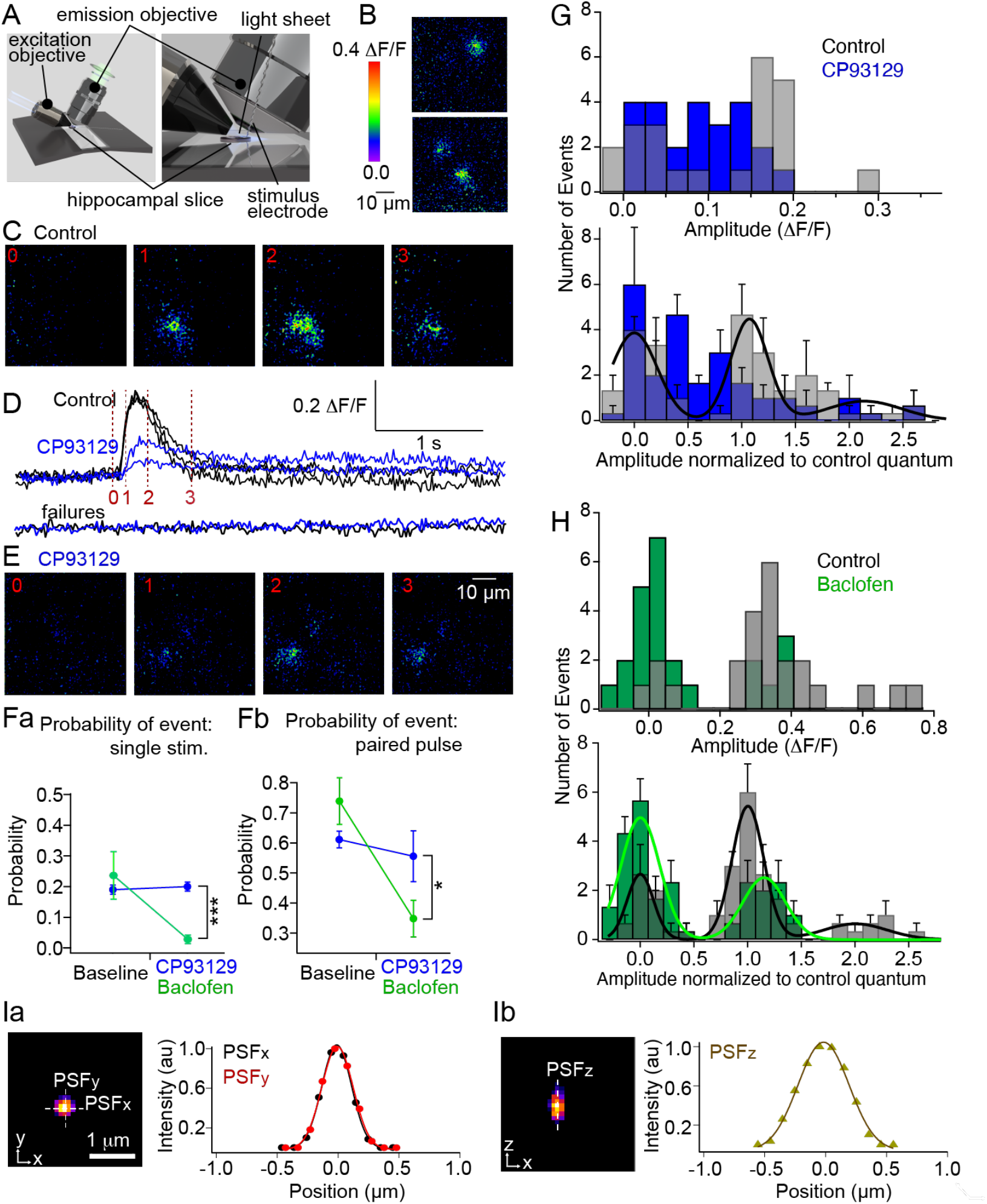
Imaging evoked glutamate release with iGluSnFR. A) Imaging of iGluSnFr expressed in subicular neurons was performed under LLSM adapted to allow recording and stimulation of hippocampal slices. The schematic shows the arrangement of preparation and lenses. B) While imaging labeled subicular pyramidal neurons dendrites, repeated stimulation (1 shock at 5 s intervals) caused stochastic fluorescent transients in subiculum. Shown are 3 responses from 2 separate stimuli. C) Evoked transients recorded in one z plane at imaged at 80 Hz expanded from a spot to larger diameter (5 – 10 µm diameter) objects (time point noted in D, red). D) Transient amplitudes and timecourses were calculated from a 5 µm diameter ROI over the transients. Shown are three control responses at the same location, obtained from 14 stimuli (black). Application of CP93129 (100 nM) reduced the amplitude of the transients that were recorded. Two responses such were recorded in the 14 stimuli in this case (blue). Examples of a failure to evoke a response in control conditions (black) and similarly in CP93129 (100 nm) (blue). E). Example of evoked transient (same location as (C) recorded in CP93129 (100 nM). Fa) CP93129 (100 nM, blue) caused no change in the probability of recording events at the same location over 21 stimuli. In contrast, baclofen (1 µM, green) significantly reduced the probability of recording and event. A two way ANOVA revealed significance between effects of baclofen and CP93129 (F(1,16) = 16.1, p = 0.001). Events were determined to be failures if the amplitude after stimulation at that location did not exceed noise. Fb) During paired pulses overall probability of recording events rose. CP93129 (100 nM, blue) again caused no reduction in probility, whereas baclofen (1 µM, green) did. A two way ANOVA revealed significance between effects of baclofen and CP93129 (F(1,32) = 7.12, p = 0.011). G) All evoked event amplitudes were measured and displayed as event/frequency histograms. Under baseline conditions events were distributed into distinct amplitudes (grey). CP93129 (100 nM, blue) removed this distinction. Top histogram (example), bottom – mean results from 5 locations normalized to the first baseline event amplitude distinct from noise. Gaussian curves were fitted to baseline data with the constraint that the amplitude and width of non-zero peaks scaled with the number of peaks. This showed a difference in control data between no response and one quantal size. Gaussian fit to the data in CP93129 was not possible. H) Similar experiment with baclofen – top histogram example, bottom average of 5 locations, before and after baclofen (1 µM, green). Baclofen left recorded amplitude events unaffected but reduced their probability. Mean data in both control (black) and baclofen (green) were fitted with gaussians with the same constraints as (G). Gaussian fits showing similar quantal peak amplitudes in both conditions. I) Point spread functions (PSFs) of the microscope obtained from a 100 nm diameter fluorescent bead. (Ia) x/y PSF, (Ib) x/z PSF of the same bead. The bead was embedded 100 µm deep in 1% agarose.

We tested the effect of 5-HT_1B_ receptor activation on these evoked fluorescence transients by recording events at single sites during repeated stimuli before and after application of CP93129 (100 nm). Frequency of events was calculated by whether an event was recorded on stimulation or not. The probability of recording an event at the same location as in control was unchanged in CP93129 0.20 ± 0.02 vs 0.19 ± 0.03 in control (Fig 6F; t(12)=0.17, p=0.87). Similar recordings were made before and during baclofen (1µM) application. In these recordings the probability of recording events was reduced significantly from 0.29 ± 0.07 in control to 0.08 ± 0.05 in baclofen (t(8)=4.24, p=0.0014).

We also measured effects of agonist application on event amplitude. However, event frequencies on single stimuli were low. To increase event frequency, stimuli were applied as paired pulses at 100 ms intervals and all post stimulation peak fluorescence transient amplitudes were determined as ΔF/F calculated from the ratio of peak response to amplitude immediately before the stimulus after background subtraction of fluorescence adjacent to the region of interest. Data was captured for each of every 24 stimuli at 5 release locations located in 3 slices for application of CP93129 and for 5 locations in 4 slices for baclofen. Event amplitudes were were plotted as frequency histograms. An example is given for baseline and in CP93129 (Fig 6G, top). In baseline conditions events largely fell into either failures, with mean signal similar to background noise, or into a signal centred on a recurring amplitude (quantum). Some larger events were also recorded. CP93129 (100 nM, blue) caused this dual amplitude distribution to be lost. Mean effects of CP93129 were compared for all 5 locations with amplitudes normalized to the mean value of a single quantum in baseline conditions (Fig 6G, bottom). The distinction between failures and a single quantum value was lost in CP93129, but there was no significant change in the number of observed failures (t(4)=1.2, p=0.15). This is emphasized by plotting a multiple gaussian curve to the baseline data with the constraints that sequential quantal events amplitudes and widths increased as integer multiples of the quantal amplitude.

Similar experiments were performed in baseline then baclofen (Fig 6H, green). In contrast, baclofen (1 µM) significantly increased the incidence of failures (t(4)=5.28, p=0.003), but remaining event amplitudes fell into the same amplitude bins as baseline responses. This is emphasized by multiple gaussian fits to baseline results and from events in baclofen in which the single event amplitude calculated from these fits was not significantly altered (t(4)=1.7, p = 0.16) whereas both the number of failures (t(4)=2.9, p = 0.04) and the number of single quantum events (t(4)=4.0, p = 0.02) were. We also measured the decay kinetics of the responses in baseline and in both agonists but no changes were measured. This is likely due to the relatively high affinities of the iGluSnFr variant used such that unbinding of glutamate from the sensor dominates the signal off rate. Indeed, the size of the events compared to the point spread function of the microscope (Fig 6I), which demonstrate submicron resolution, indicates that we detect glutamate spillover from the synapse as a large part of the signal.

### Simulating glutamate release and receptor activation

GABA_B_ receptors reduce P_r_, whereas, 5-HT_1B_ receptors lower cleft glutamate concentrations. The latter may be due to a modification of the fusion pore during neurotransmitter release (Harata et al., 2006; Photowala et al., 2006), changes in MVR (Rudolph et al., 2015) or modulation of the location of release with respect to postsynaptic receptors (Tang et al., 2016; Chen et al., 2018; Haas et al., 2018). We created a Monte Carlo simulation of a simple synapse using MCell to probe these possibilities.

The simulated synapse comprised a disk-shaped synaptic cleft, 300 nm in diameter and 20 nm deep, with kinetically modeled AMPA (Robert and Howe, 2003) and NMDA receptors (Banke and Traynelis, 2003) placed on the postsynaptic surface (Fig 7). For each condition, 20 random seeds were simulated, whereby glutamate release was modeled from between one to three vesicles attached to the presynaptic surface with a dynamically opening pore (Multimedia Video 1), initially with a 0.4 nm diameter and the length of two lipid bilayers. This is approximately the value proposed for the conductance of foot events in large dense core vesicles (Vardjan et al., 2007). The simulated pore then expanded to the full diameter of the vesicle over a range of timecourses from 0.2 to 20 ms. We tested three hypotheses: 1) glutamate release was slowed through a fusion pore, 2) multivesicular vesicle numbers were reduced, 3) relative locations of release sites and receptors were altered. The parameter files for these simulations are available on our website (https://alford.lab.uic.edu/GPCRs.html). The individual parameters and sources are listed in Table 1.

**Figure 7.**
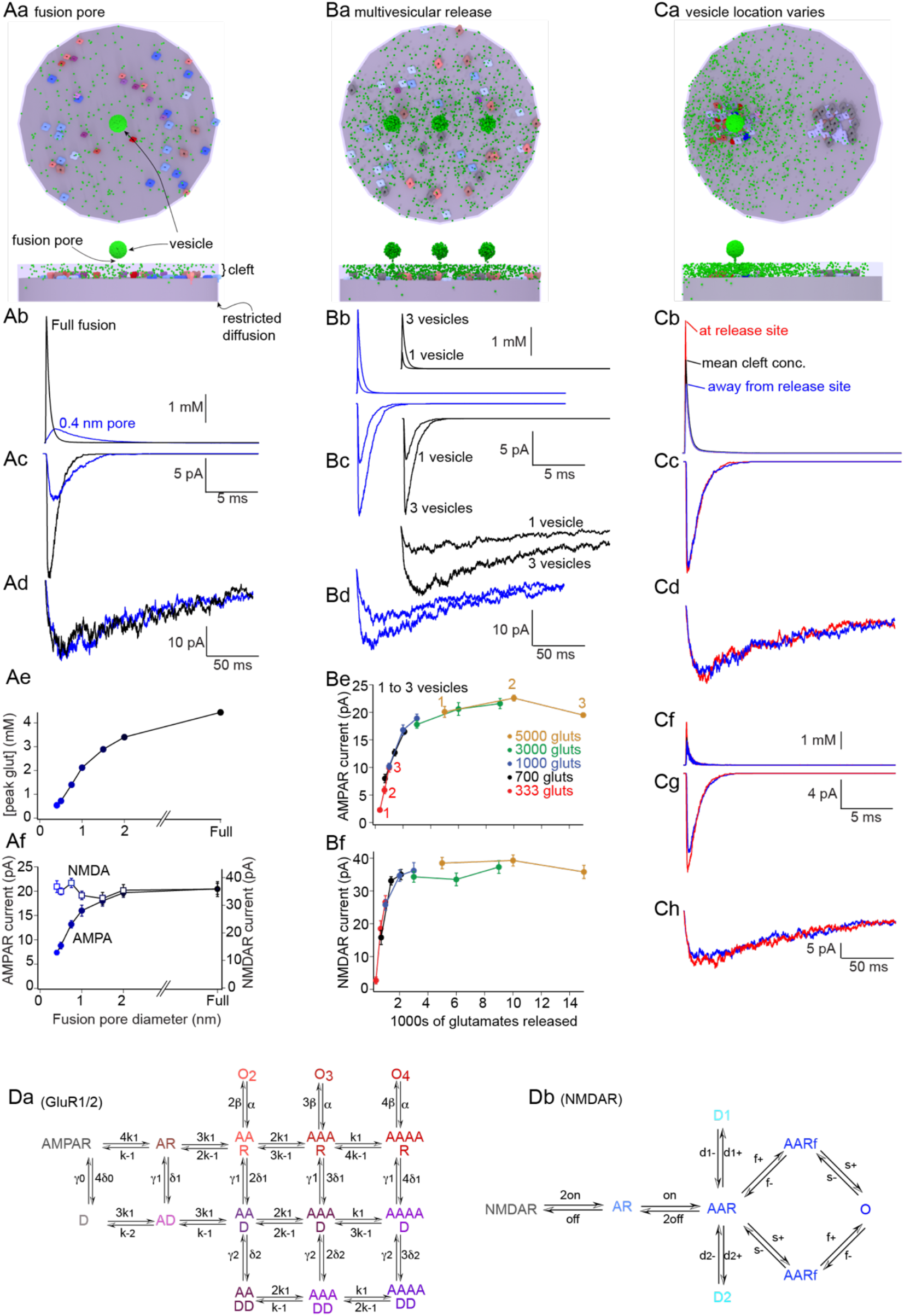
Simulating neurotransmitter release and receptor activation. Aa) Simulated 300 nm diameter synapses, 20 nm deep synaptic clefts and a 1.7 nm path at the edge to restrict fusion. Image is from 500 µs after the start of fusion. To simulate a fusion pore, fusion started from a pore at fixed diameter (0.4 nm for 20 ms) before fully opening (Aa) or pores that fully opened within 200 µs. Simulated receptors were randomly seeded on the postsynaptic surface (receptor colors as for kinetic models (D). Ab) Varying initial fixed pore diameters slowed glutamate release, reduced its cleft concentration (graphed Ae) and differentially inhibited AMPA (Ac) over NMDA (Ad) receptor-mediated responses (graphed Af, AMPA filled circles, NMDA hollow squares). Ba) Simulated multivesicular release pattern. 3 simulated vesicles where vesicle number was varied from 3 to 1 with 333 to 5000 glutamates/vesicle. Bb) The effect on cleft glutamate concentration of reducing vesicle numbers from 3 to 1 when each contained either 1000 glutamates (blue) or 700 glutamates (black). The AMPA receptor response (Bc) and the NMDA receptor responses (Bd) are shown below. Graphs show the AMPA (Be) and the NMDA receptor responses (Bf). To each condition, both varying glutamates/vesicle (color coded) and number of vesicles, each point in the colored sections. Ca) The effect of varying the position of the release site with respect to postsynaptic clustered receptors was simulated. Two receptor clusters were seeded with a mean separation of 150 nm and the releasing vesicle positioned over one. Cb) cleft glutamate concentrations over the clusters (red at release site, blue away from release) and mean cleft concentration (black) after 5000 glutamates were released in 200 µs. Cc) AMPA receptor responses simulated under the release site (red) and away from the site (blue). Cd) NMDA receptor responses. Cf-h) Similar experiment to Cb-d, but with 1500 glutamate molecules to reveal modulation of AMPA and NMDA receptor mediated responses. Da) Kinetic model for AMPA and (Db) for NMDA receptors (parameters in table 1). Colors are the same as shown for receptors in Aa, Ba and Ca.

To simulate a slowly opening fusion pore, the vesicle (containing 5000 glutamates) was attached to the presynaptic membrane through a pore (Fig. 7A). The simulated pore expanded to the diameter of the vesicle, but arrested at a diameter of 0.4, 0.5, 0.75, 1.0, 1.5 or 2nm for 20ms. Receptors were randomly seeded on the postsynaptic membrane, and their states represented by colors in kinetic models (Fig. 7D). Arresting the fusion pore opening at set diameters slowed release and reduced peak cleft glutamate concentrations (Fig. 7Ab) compared to full expansion of the pore to the diameter of the vesicle within 200 µs, which represents the default state. This slower glutamate release reduced simulated AMPA, but not NMDA receptor responses (Fig.7Ac-Ad) (Multimedia Video 2). Both the peak simulated glutamate concentration and peak AMPA receptor activation were reduced with smaller pore sizes, whereas NMDA receptor responses remained unchanged (Fig 7. Ae-Af).

MVR was simulated by varying numbers of simulated fusing vesicles from 1 to 3 (Fig. 7B). One vesicle was placed at the center and two offset by 75nm (Fig. 7Ba). Fusion was simulated by pores fully opening in 200 µs. Vesicle content varied from 333 to 5000 glutamates because at high glutamate content the AMPA receptor response saturated with more than one vesicle, preventing any modulation. Varying vesicle numbers altered cleft glutamate concentration and modulated AMPA and NMDA receptor responses nearly equally (Fig. 7Bb-Bd). The effect on both NMDA receptors and AMPA receptors was modulated by the total number of released glutamates rather than vesicle number (Fig 7Be-Bf).

To vary vesicle fusion position compared to receptors, receptors were seeded within two proscribed areas of the postsynaptic density; one under the fusing vesicle and the other a mean distance of 150 nm away (Fig. 7Ca). Release of 5000 glutamates was simulated with complete fusion in 200 µs causing peak glutamate concentrations that differed at the two sites. Little modulation of AMPA or NMDA receptor responses occurred (Fig. 7Cc-Cd). Earlier work simulating vesicles displaced from the release site reported larger effects with fewer released glutamates (Haas et al., 2018). Consequently, the simulation was repeated with 1500 glutamates (Fig. 7Cf-Ch). The more distant clusters of AMPA and NMDA receptor responses were reduced to 80% and 83% of the amplitude under the release site respectively. Thus, the response is sensitive to relative locations of vesicle fusion and receptor location, but this effect is small compared to that of experimental 5-HT_1B_ or GABA_B_ receptor activation and does not recapitulate the difference in effect between AMPA and NMDA receptor responses. Overall, changing fusion pore dilation best mimicked the physiological effect of 5-HT_1B_ receptor activation.

### Simulating the amplification of inhibition by kynurenate

Experimental effects of the low affinity antagonist, kynurenate, on AMPA receptor EPSCs indicate that 5-HT_1B_ receptors, but not GABA_B_ receptors, lower the concentration of glutamate in the synaptic cleft. To determine whether kynurenate might discriminate between mechanisms by which cleft glutamate concentration is changed based on receptor effects, we simulated the action of kynurenate using a modification of the kinetic model of the AMPA receptor.

The kinetic model assumed that kynurenate can substitute for glutamate at all states of the receptor but prevented receptor channel opening (Fig. 8Aa). Simulations were performed with kynurenate (240 µM) present in the synaptic cleft. This reduced the simulated AMPA receptor-mediated response to 55% of control (Fig. 8Ab). The simulations were then repeated with the synaptic vesicle fusion pore diameter arrested for 20 ms at diameters from 0.4 to 1.5 nm. The effect of kynurenate was enhanced when the pore size was restricted (Fig. 8B). The effect of simulated kynurenate was also tested against MVR. Modifying the number of fused vesicles was only effective at a reduced number of glutamates present in each vesicle, thus, these simulations were performed from with three, two or one vesicles each containing 700 glutamates as for Fig 7Bb-d. Kynurenate also enhanced the effect of reduced vesicle number on the simulated AMPA receptor response (Fig. 8C).

**Figure 8.**
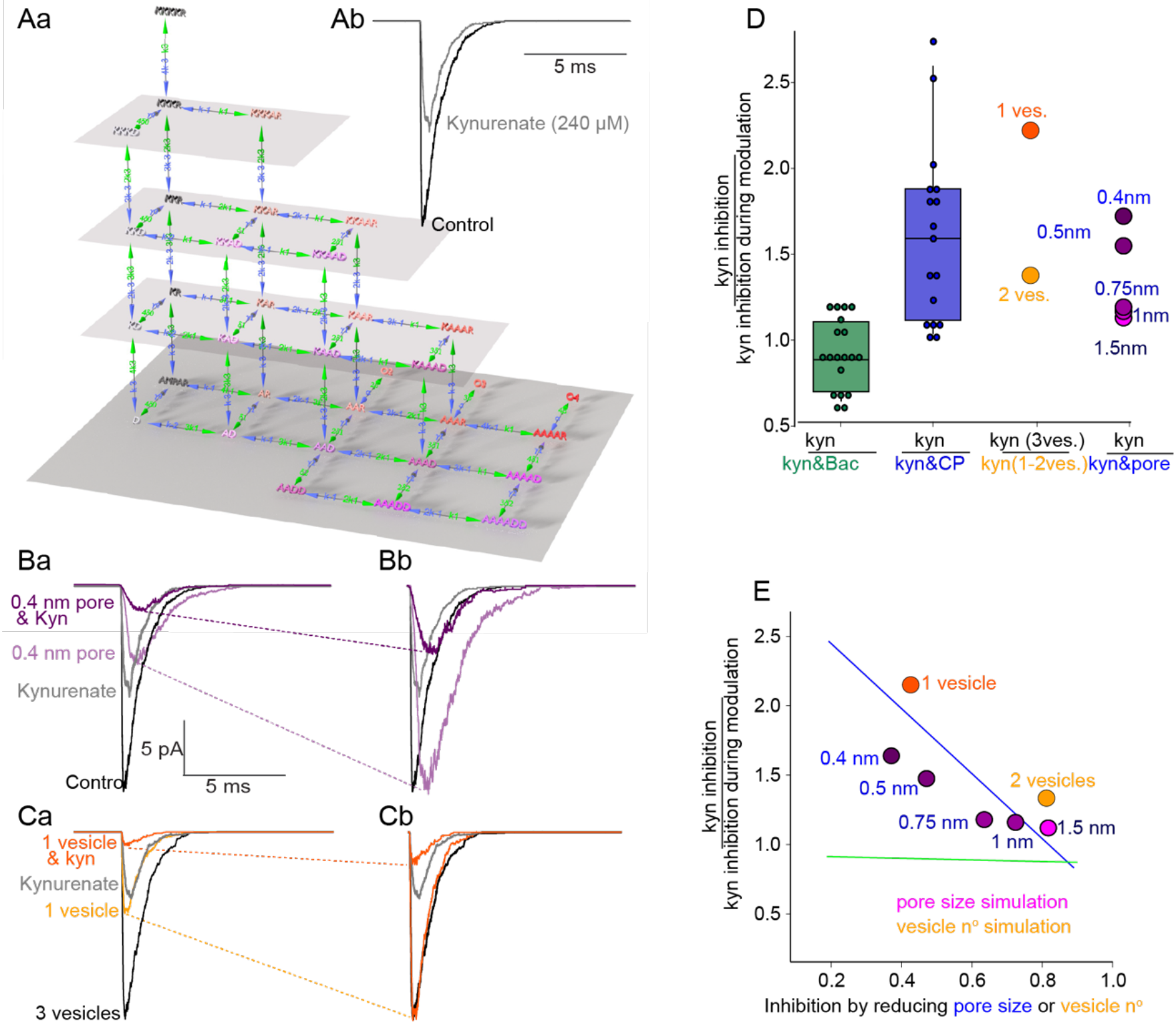
Simulating cleft glutamate modulation and kynurenate inhibition. Aa) Kinetic model of AMPA receptor activation by glutamate and block by kynurenate. (Colors and lowest tier of 3D model are as for Fig 7Da, Vertical elements of the 3D representation show kynurenate occupying receptor bound and desensitized states, parameters in table1). Ab) Simulated full fusion in 200 µs with 5000 glutamates (black, control) and the effect of 240 µM kynurenate (gray). Ba) The effect of kynurenate (gray) on control simulated responses (black) and on the response when the vesicle fusion pore was restricted to 0.4 nm (light purple without, purple with kynurenate). Bb) Full fusion responses and 0.4 nm pore responses scaled to emphasize effect of kynurenate is amplified by the 0.4 nm pore. Ca) Simulation as numbers of vesicle were varied from 3 to 1, each with 700 glutamates as in Fig 7Bb-d. Effect of kynurenate (gray) on 3 vesicle response (black), and on 1 vesicle response (light orange) plus kynurenate (orange). Cb) Responses scaled to compare effect of kynurate on the 3 vesicle and 1 vesicle responses. D) Comparison of magnification of kynurenate inhibition by baclofen (green) and CP93129 (blue) from experimental data (Fig 3E) to the effect on kynurenate inhibition in simulations of reducing vesicle numbers (orange) or changing the initial fixed vesicle pore diameter (purple). E) Simulation data (from D) plotted against the effect of reducing vesicle number (orange) or fusion pore size (purple) on AMPA receptor responses. The green line is the fit to the similar effect of kynurenate on variations in experimental baclofen inhibition (from Fig 3G). The blue line similarly for experimental CP93129 variation (from Fig 3H).

Effects of both these manipulations were compared to the experimental effects of GABA_B_ and 5-HT_1B_ receptor activation on AMPA mediated synaptic responses. Box plots (from Fig. 3E) showing CP93129 and baclofen effects on kynurenate mediated antagonism are presented alongside simulated kynurenate inhibition following a change in number of fusing vesicles (normalized to the maximum – 3) and following arrest of the fusion pore at between 0.4 and 1.5 nm diameters normalized to full fusion (defined as complete opening of the fusion pore the vesicle diameter in 200 µs). Both of these manipulations enhanced the efficacy of kynurenate (Fig. 8D).

As for the experimental effects of agonists (Fig 3E,H), simulations of variation in inhibition of the AMPA response, either by varying the number of multivesicular fusing vesicles at one active zone (Fig. 8D orange) or varying the vesicle pore diameter (Fig. 8D purple) correlated with the enhancing effect of kynurenate on both manipulations (Fig. 8E). Both approaches recapitulated experimental effects of CP93129 but not of baclofen (Fig. 8E). From this combination of experimental effects of low affinity antagonist, and its simulation, the experimental data can be explained by a reduction synaptic cleft glutamate. Results with kynurenate alone, however, do not discriminate between the effect on release, whether change in fusion pore diameter or change in fusing vesicle numbers that causes this change in concentration.

### Modulation of the timecourse of the AMPA receptor mediated response

Simulating the slowing of the fusion pore opening slowed the rise and decay times of the simulated AMPA receptor response (see Fig. 7Ac) demonstrated by replotting the inhibited response scaled to the peak of the control (Fig. 9A). In simulated data, the arrested pore diameter (0.4 nm) increased the timecourse (τ) of the rise from 0.09 to 0.35 ms and of the decay from 0.97 to 1.59 ms measured by single exponentials fitted to the rise and decay. Simulated kinetic properties of AMPA receptor opening are more rapid than those experimentally obtained. This may be because they are neither affected by dendritic filtering nor by asynchronous vesicle fusion. Nevertherless, we investigated effects of agonist on the kinetics of synaptic responses.

**Figure 9.**
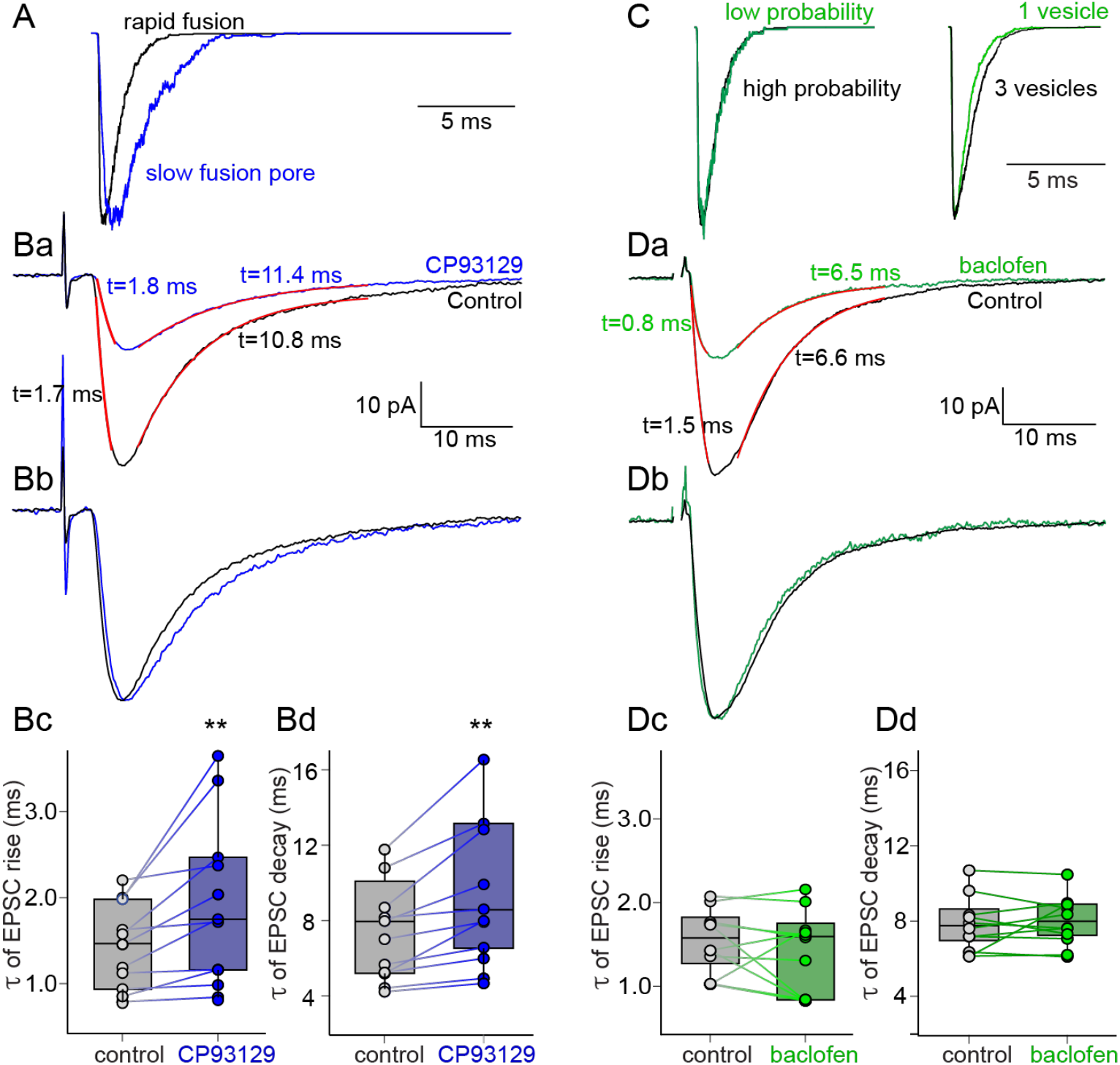
Effect of agonist on AMPA-mediated EPSC kinetics. A) Data from simulating full fusion and a 0.4 nm pore scaled to the same amplitude to show the slowing of the rise and decay times of the resultant AMPA receptor-mediated response. Ba) Experimental data of control responses (mean of 10 sequential EPSCs, black) and responses in the same neuron in CP93129 (50 nM, blue). Red curves are single exponentials fitted to 10 to 90% of the rise and of the decay phases. Bb) response amplitudes scaled to emphasize the change in rates of the response in CP93129. Bc) Box plots and data points for the risetime in control (gray) and in CP93129 (blue), Bd) for decay times. C) Simulated AMPA mediated EPSCs scaled to show the kinetic effect of reduced number of responses to mimick a change of P_r_ at univesicular synapses on simulated kinetics (right) or reduced number of vesicles (from 3 to 1) at multivesicular synapses (left). Da) Experimental control responses (black) and in the same neuron in baclofen (600 nM, green). Red curves show single exponentials fitted to rise and decay phases as in B. Db) Response amplitudes scaled as in B. Dc) Box plots and data points for the risetime in control (gray) and in baclofen (green), Dd) similarly for decay times.

Responses from 11 neurons were recorded in which EPSCs showed monotonic decays (Fig. 9B). CP93129 (50nM) increased τ’s of rise and decay phases, visible when the response in CP93129 was scaled to the control amplitude (Fig. 9Bb). Single exponential fits to the 10 to 90% risetimes showed a significant increase from a mean of 1.4 to 2.0 ms (t(10) =3.06, p=0.006) as did the decay times from 7.3 to 9.4ms (t(10)= 3.54, p=0.003) (Fig.9B).

Simulations of reduction of P_r_ between synapses is trivial – a reduction in numbers of sites – consequently kinetic properties of the simulated EPSC are unchanged (Fig. 9C, left panel). Alternatively, a simulated reduction in P_r_ at a multivesicular synapse shows a slight decrease in decay rate (Fig. 9C, right hand panel). We compared this to effects of baclofen (1µM), which reduces P_r_, in 10 neurons, again in which the decay was monotonic. Baclofen had no significant effect on mean rise time (control=1.56 ± 0.12ms, baclofen=1.44 ± 0.16 ms; t(9)= 0.83, p=0.21) or decay (control=7.92 ± 0.47ms, baclofen=8.11 ± 0.41ms; t(9)= 0.59, p=0.30) (Fig.9D). These results emphasize that 5-HT_1B_ and GABA_B_ receptors on the same presynaptic terminals mediate inhibition by different mechanisms.

### Repetitive stimulation and train-dependent responses

At the same terminals, GABA_B_ receptors inhibit Ca^2+^ entry, whereas 5-HT_1B_ receptors cause Gβγ to interfere with Ca^2+^-synaptotagmin SNARE complex interactions (Hamid et al., 2014). This implies selective effects of receptor activation, but also synergy following activation of both receptors (Zurawski et al., 2019). As presynaptic Ca^2+^ accumulates during stimulus trains (Yoon et al., 2007; Hamid et al., 2014) that induce long term plasticity (O’Keefe, 1990), it may modify competition for interaction with the SNARE complex between Gβγ and synaptotagmin or other Ca^2+^ sensors such as Doc2 (Groffen et al., 2006) (Groffen et al., 2006) or synaptotagmin 7 (Jackman et al., 2016; Jackman and Regehr, 2017).

GABA_B_ receptors inhibit train dependent Ca^2+^ accumulation. To show this, we recorded Ca^2+^ transients in CA1 pyramidal neuron terminals by filling the axons with Ca^2+^ sensitive dyes. CA1 neurons were whole cell recorded under current clamp with a red dye (Alexa 594) to determine the axon location and a Ca^2+^ sensitive dye (Fluo 5F) to record action potential evoked Ca^2+^ transients. We have previously shown that Fluo 5F allows reliable recording of Ca^2+^ signals (Hamid et al., 2014; Hamid et al., 2019). Individual presynaptic varicosities in the subiculum were imaged using line scanning (500 Hz) and neurons stimulated with 2 ms depolarizing current pulse trains (5 stimuli at 20 ms intervals) to evoke action potentials (Fig 10A, upper trace). Line scanning resolved Ca^2+^ transients over accumulating Ca^2+^ during the stimulus train (Fig 10Aa, lower trace). Baclofen (600 nM) reduced the Ca^2+^ transient following each stimulus in the train (Fig 10Ac, green squares). The last transient was reduced to 0.46 ± 0.07 of control (t(3)=2.94, p=0.049). CP93129 (100 nM), had no effect on the train evoked Ca^2+^ transients (Fig 10Ac, blue circles). The last transient in the train in CP9319 was 1.18 ± 0.11 of control (t(3)=1.55, p=0.13). The absolute value of Ca^2+^ surrounding synapses may be lower than used in these recordings (Borst, 2010), although the values of P_r_ that we obtained from iGluSnFr experiments were similar, but the overall finding that 5-HT_1B_ receptor effects are both Ca^2+^ and train dependent holds true.

**Figure 10.**
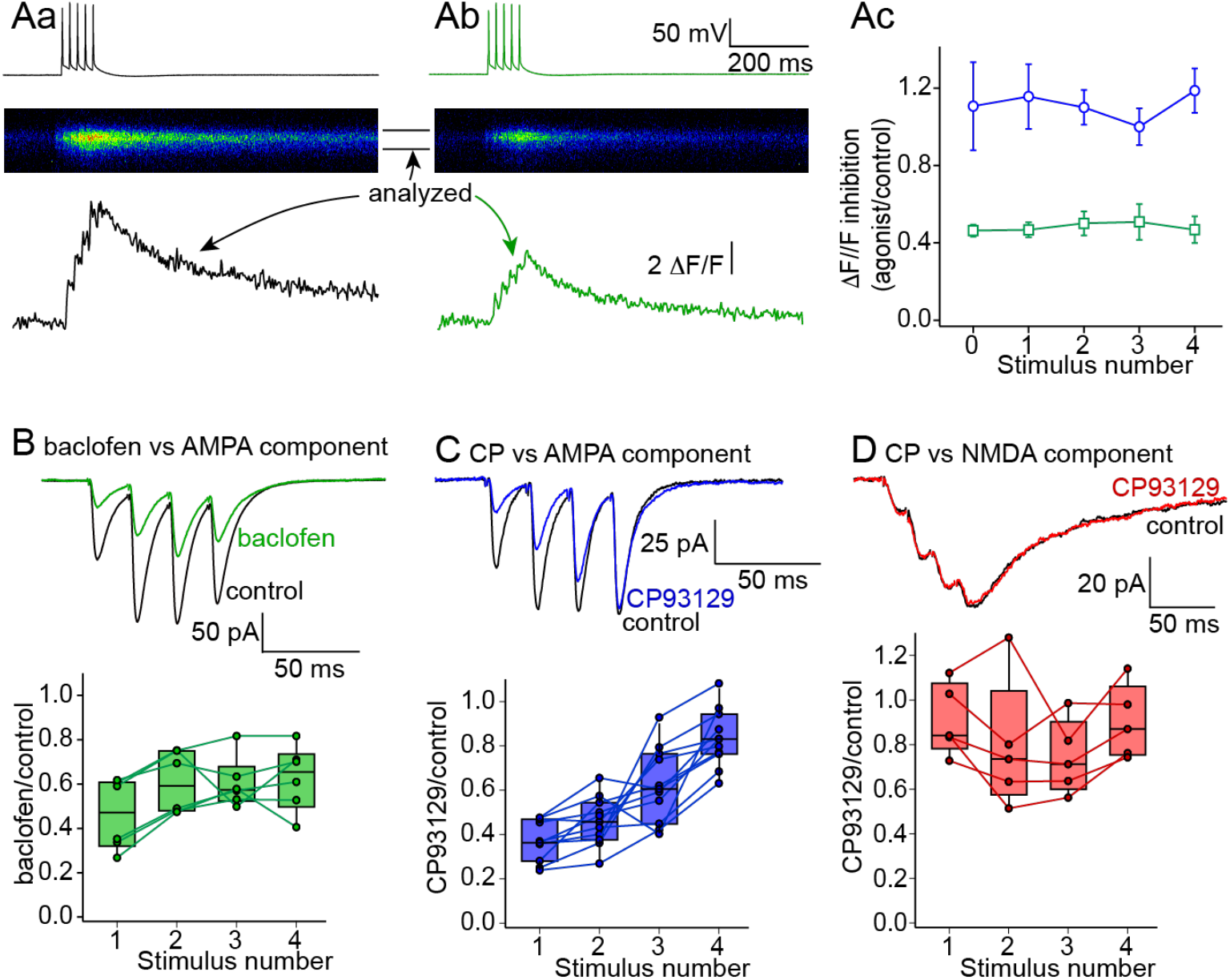
Effect of repetitive stimulation on inhibition. A) Line scans through single CA1 neuron presynaptic varicosities imaged from neurons filled with Ca^2+^ dye (Fluo5F) from whole cell patch recordings. Aa) A train of 5 action potentials (top) were triggered via the patch pipette to evoke a train of fluorescence transients in the varicosity (middle) that were quantified (bottom). Ab) Baclofen (600nM) did not prevent evoked events but reduced the Ca^2+^ transient amplitude following each action potential. Ac) Means of peak amplitudes in baclofen divided by pre-agonist control (green squares) of the peak of each transient in the train, from 4 varicosities. In contrast in a further 4 varicosities, CP93129 (1 µM, blue circles) had no effect on transient amplitudes throughout trains. B) Train of stimuli (4 shocks, 50Hz) applied to CA1 axons to evoke AMPA EPSCs recorded in subicular neurons (in D-AP5 and bicuculline at a holding potential of −60mV). Baclofen (1 µM) inhibited EPSCs throughout the train of stimuli. C) A similar train stimulus applied for isolated AMPA receptor mediated EPSCs in which Box plots and raw data are as for (B). CP93129 (400nM, blue) inhibited the first response in the train, but the EPSCs recovered to control amplitudes by the fourth stimulus. D) A similar train stimulus applied in a preparation superfused with NBQX and bicuculline to isolate the NMDA receptor EPSCs. EPSCs were recorded at −35mV to eliminate Mg^2+^ block of the NMDA receptor. CP93129 (400nM) had little effect on any of the responses in the stimulus train. Box plots and raw data as for B.

Consequently, we hypothesized that modifying Ca^2+^ entry with GABA_B_ receptors will alter the efficacy of 5-HT_1B_ receptors during repetitive activity to provide a form of metamodulation and synaptic integration. To address this, we first examined effects of stimulus trains (4 shocks, 50Hz) on inhibition of synaptic responses by CP93129 and baclofen. AMPA EPSCs were isolated in bicuculline and D-AP5. Baclofen inhibited AMPA EPSCs (measured from EPSC initial slopes) throughout these trains with no significant difference in inhibition between responses within the train (other than the paired pulse enhancement, Fig 2) (Fig. 10B; ANOVA single factor F(3,20)=28.2, p=0.18). In CP93129 (100nM) initial slopes of the 1^st^ responses were reduced to 42±3% of control, while the 4^th^ only to 78±9% (Fig. 10C; single factor ANOVA determined whether CP93129 differentially altered EPSCs in the train - this was significant, ANOVA single factor F(3,40)=28.2, p=5.7 x 10^−10^). Post hoc Tukey HSD analysis demonstrated the 3^rd^ and 4^th^ EPSCs were inhibited significantly less than the 1^st^ (p<0.001). No significant inhibition of NMDA EPSCs isolated in bicuculine (5µM) and NBQX (5µM) was recorded for any stimulus (Fig. 10D; ANOVA single factor F(3,16)=0.36, p=0.78).

### GPCRs filter the final output of the hippocampus

GABA_B_ receptors provide fairly uniform inhibition of synaptic output during stimulus trains after the second stimulus (Fig. 10B), when paired pulse facilitation associated with changes in P_r_ is increased. By contrast, 5-HT_1B_ receptor activation reveals a complex modification of synaptic outputs from the hippocampus. NMDA receptor-mediated EPSCs are barely effected, though *in-vivo* activation of the NMDA responses would require depolarization, which can be provided by synaptic transmission via AMPA receptors. AMPA receptor responses are blocked at the start but not the end of short stimulus trains. If 5-HT_1B_ receptor activation merely inhibited AMPA receptor responses then synaptically evoked depolarization would be blocked and NMDA receptors would remain blocked by Mg^2+^. Therefore, we first determined how 5-HT_1B_ receptors modify information flow between CA1 and subicular neurons during activity trains in which AMPA receptor-mediated depolarization triggers NMDA receptor-mediated responses (Herron et al., 1986).

We stimulated CA1 axons with short stimulus trains as earlier but recorded from subicular neurons under current clamp conditions. Stimulation intensity was adjusted (up to 20 μA) to ensure that the first stimulus of the burst invariably caused an action potential in the recorded postsynaptic cell (Fig 11Aa). We found a complex mode of inhibition, mediated by CP93129. After application of the agonist, the first EPSP is significantly inhibited, leading to a dose-dependent delay in depolarization and consequent firing of an action potential (Fig 11A-D), but CP93129 did not prevent action potential firing during the train, even at saturating concentrations. The mean number of APs following the first stimulus in 4 cells was 1.3±0.2 in control, but reduced to 0.2±0.1 by a half maximal concentration of CP93129 (50nM, p<0.05, Fig.11Ea). However, firing recovered by the 2^nd^ stimulus, in which the control stimulus evoked 1.1±0.4 APs and stimuli in CP93129, evoked 1.1±0.4 (Fig.11Eb), while by the 3^rd^ stimulus, evoked APs were enhanced by CP93129 (from 0.4±0.2 to 0.7±0.3, increase in number from1^st^ to 3^rd^ was significant, p<0.01). Thus, while we see a concentration-dependent CP93129-evoked delay of APs (Fig.11A-D). The mean area under the depolarization is barely altered (86±5% of control at 1µM), even when the first AMPA receptor-mediated response is substantially inhibited (to 10±2%, Fig.11F). This reflects the frequency-dependent recovery from 5-HT_1B_ receptor-mediated inhibition and lack of a CP93129 effect on NMDA receptor-mediated EPSCs.

**Figure 11.**
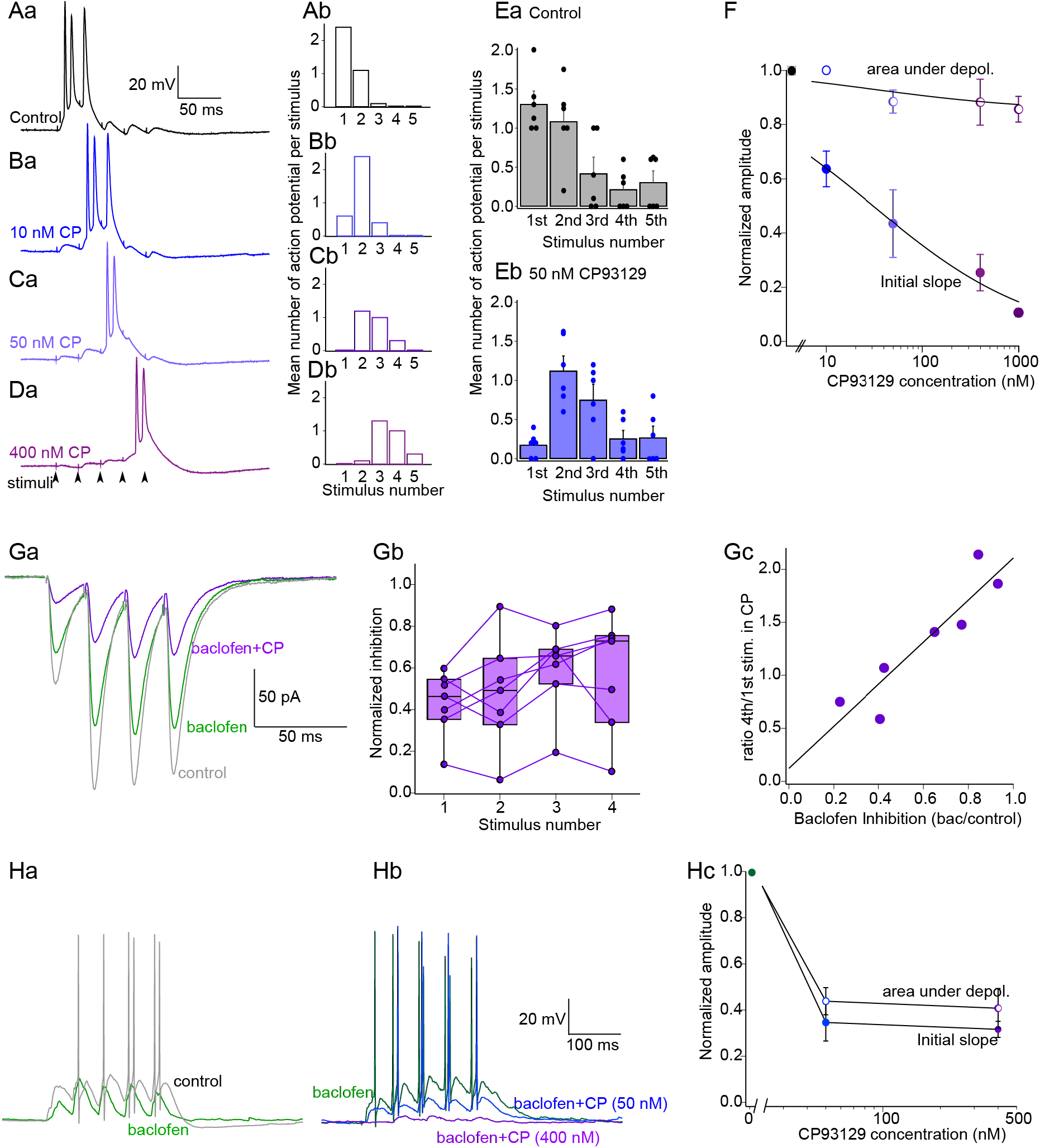
5-HT_1B_ receptors filter synaptic output of the hippocampus but these filter properties are modified by GABA_B_ receptors. A) 5-HT_1B_ receptors suppress activity at the start of a train but not total activity. A stimulus train (5 shocks, 50 Hz) was applied to CA1 axons, EPSPs recorded from subicular neurons. Stimulus intensity was adjusted to ensure the first train stimulus caused an AP. (Aa) Example control trace. (Aa) Histogram shows mean number of APs following each stimulus from 10 sequential trains from this cell. (Ba-Da) Recordings of the cell in (A) after in increasing concentrations of CP3129 (CP). (Bb-Db) AP counts from each stimulus in the recordings. E) Mean AP counts from each stimulus from 4 cells. (Ea) in control and (Eb) in 50nM CP93129. F) CP93129 inhibits the first EPSP but not total excitation during trains. From 4 cells a comparison of dose dependence of CP93129 inhibition of the mean EPSP initial slope on stimulus 1 and mean area under the depolarization for all stimuli (data normalized to control). G) GABA_B_ receptor activation modifies the filtering effect of 5-HT_1B_ receptors. Ga) As shown in Fig 10, baclofen (green) 1 µM inhibits EPSPs uniformly throughout the stimulus train. CP93129 (400 nM purple) in the presence of baclofen now also inhibits uniformly throughout the train. Examples are mean responses from 10 sequential trains in one cell. Gb) Box plots and single data show effects in 7 neurons of adding CP93129 (400nM) to baclofen on responses from each of the 4 stimuli in the train. Gc) the effect of CP93129 on the last EPSC in the train is proportional to the initial inhibtory effect of baclofen. Ha) The modulating effect of Baclofen 1 µM on CP93129 inhibition is seen in current clamp recordings. Baclofen (green) inhibited EPSPs throughout the train. Hb) In the same recording in baclofen the stimulation intensity was increased (green). CP93129 was added at 50 nM (light blue) and 400 nM (dark blue). EPSPs in baclofen were now uniformly inhibited throughout the stimulus train. Hc) graph of amplitudes normalized to response in 1 µM baclofen for the first EPSP in the train and for the total area under the depolarization in increasing CP3129 concentrations.

Baclofen inhibits EPSCs throughout trains (Fig.10B), and have previously been shown to act by inhibiting Ca^2+^ entry (Hamid et al., 2014). This implies it can modify Ca^2+^-synaptotagmin competition with Gβγ at SNARE complexes and consequently meta-modulate effects in CP93129 which causes Gβγ competition with Ca-synaptotagmin at SNARE complexes (Yoon et al., 2007). We investigated combined receptor activation on presynaptic modulation. Baclofen (1µM) Inhibited AMPA receptor EPSCs uniformly during stimulus trains (Fig.11Ga - green). However, in baclofen, CP93129 now also inhibits EPSCs throughout the train (Fig.11Ga,Gb - purple). This meta-modulation of CP93129 inhibition by baclofen depends on the initial efficacy of baclofen. Plotting initial effects of baclofen alone against CP93129 inhibition of the last EPSC in the train in baclofen, reveals a linear relationship (Fig.11Gc). This is consistent with GABA_B_ receptor inhibition of Ca^2+^ channels reducing Ca^2+^ that causes evoked release at each stimulus and for Ca^2+^-synaptotagmin competition with 5-HT_1B_ receptor released Gβγ at SNARE complexes.

We tested this effect of baclofen on CP93129 filtering of EPSP trains. We did not evaluate specific effects of baclofen alone because polysynaptic inhibition is also altered by baclofen and these current clamp experiments included no GABA_A_ antagonists. Nevertheless, recordings in baclofen provided a starting condition to test effects of CP93129 during trains after partial inhibition of presynaptic Ca^2+^ entry by baclofen (Fig. 11 Ha). CP93129 in baclofen reduced the area under the depolarization throughout the train (Fig. 11Hb), such that in contrast to the effects of CP93129 alone, in baclofen CP93129 equally inhibited the first EPSP slope and the area under the depolarization during the train. Changes in area and initial slope are not significantly different at 50 or 400nM CP93129 (Fig. 11H t(7)=1.45, p=0.19 and t(3)=1.39, p=0.252 respectively). Thus, different receptors with different effectors inhibit release with different temporal outcomes through train evoked responses, but they also interact such that one receptor (GABA_B_) alters the effect of a second (5-HT_1B_) both qualitatively and quantitatively causing an entirely novel form of presynaptic integration.

## Discussion

Two well-characterized pathways mediate GPCR membrane-delimited presynaptic inhibition. Gβγ inhibits presynaptic Ca^2+^ entry or competes with Ca^2+^-syt1 at the SNAP-25 c-terminal. GABA_B_ receptors cause presynaptic inhibition throughout the CNS (Dutar and Nicoll, 1988; Alford and Grillner, 1991). They are thought to act principally via Gβγ inhibition of Ca^2+^ channels (Wu and Saggau, 1995; Hamid et al., 2014), although, in the nucleus accumbens, they target SNARE complexes (Manz et al., 2019). Understanding their function is important given use of baclofen in disorders such as multiple sclerosis, cerebral palsy, spinal injury and stroke. Serotonin also modulates central excitability, and 5HT1 receptors are found presynaptically. 5-HT_1B_ receptors control glutamate (Zifa and Fillion, 1992; Sari, 2004) and 5-HT (Davidson and Stamford, 1995) release, and play a role in delayed action of Selective Serotonin Reuptake Inhibitors (SSRI) (Svenningsson et al., 2006; Tiger et al., 2018), in anxiety and depression (Nautiyal et al., 2016). Two presynaptic targets have been found. In the Calyx of Held, they reduce Ca^2+^ entry (Mizutani et al., 2006), and In lamprey, they modify synaptotagmin-SNARE interactions (Blackmer et al., 2001; Blackmer et al., 2005; Gerachshenko et al., 2005). GABA_B_ and 5-HT_1B_ receptors colocalize at CA1 pyramidal neuron terminals (Boeijinga and Boddeke, 1993; Hamid et al., 2014) where 5-HT_1B_ receptors cause Gβγ-SNAP-25 interactions, and GABA_B_ receptors inhibit Ca^2+^ entry (Hamid et al., 2014; Zurawski et al., 2019). Why receptors colocalize to cause presynaptic inhibition via different mechanisms may depend on their intersecting signaling targets.

A third mechanism is possible, whereby neuromuscular miniature endplate potential frequency is reduced (Silinsky, 1984). Similarly a mechanism independent of Ca^2+^ entry or interaction with c-terminal SNAP-25 inhibits miniature (m)EPSCs in hippocampal neurons (Rost et al., 2011). The mechanisms that we have analyzed at CA1 terminals do not involve this effect. GABA_B_ receptors inhibit Ca^2+^ entry, precluding effects on mEPSCs. Gβγ from 5-HT_1B_ receptors competes with syt1 to reduce release, but syt1 does not evoke mEPSCs (Pang et al., 2011; Vyleta and Smith, 2011), therefore, we would not expect Gβγ to alter mEPSCs. While, Gβγ-SNARE complex interaction might alter mEPSCs, clear results are hard to obtain because inputs to subicular pyramidal neurons come from mixed populations while only CA1 terminals express 5-HT_1B_ receptors. Gβγ might alter other Ca^2+^ sensor/SNARE interactions to mediate these effects (Yoon et al., 2008), but we have no evidence of this at CA1 synapses. Furthermore, we have previously demonstrated that 5-HT_1B_ receptors do not inhibit Sr^2+^ evoked asynchronous release at CA1 terminals (Hamid et al., 2014). Sr^2+^ does not trigger fusion via syt1 SNARE interactions in reduced fusion models (Bhalla et al., 2005) indicating that asynchronous release is triggered by other fusogenic proteins – e.g. Doc2 (Yao et al., 2011).

Targets of presynaptic GPCRs determine their effects on neurotransmission. For example, Ca^2+^ channel inhibition reduces P_r_, and in CA1 terminals, GABA_B_ receptors enhance PPR, but leave synaptic glutamate concentration unaffected. They also equally inhibit NMDA and AMPA responses. In contrast, 5-HT_1B_ receptors do not change P_r_. They leave PPRs unaffected, do not modify presynaptic Ca^2+^ entry (Hamid et al., 2014), but inhibit AMPA and NMDA EPSCs differentially. Therefore, we investigated variations in cleft glutamate.

Changing P_r_ with Mg^2+^ or baclofen left synaptic glutamate concentration unaltered, whereas 5-HT_1B_ receptors lowered it. We also found that 5-HT_1B_ receptors inhibited AMPA receptor-mediated EPSCs more than NMDA EPSCs. Low affinity antagonism of NMDA and AMPA receptors shows that cleft glutamate concentrations fall during 5-HT_1B_ receptor activation. Using iGluSnFr to visualize glutamate release revealed that 5-HT_1B_ receptors reduced the amount of glutamate released, but not the probability of each event. In contrast, GABA_B_ receptor activation reduced the probability of recording an event on each stimulus. It increased the numbers of failures and reduced the number of discrete amplitudes recorded.

At least three explanations exist for these findings. (1) 5-HT reduces P_r_ where MVR predominates (Rudolph et al., 2015) and simultaneous fusion of multiple vesicles are linked (Singer et al., 2004). In this scenario a change in P_r_ by 5-HT is hidden by de-linking simultaneous fusion of multiple vesicles. (2) 5-HT modifies subsynaptic locations of release (Biederer et al., 2017; Chen et al., 2018). (3) A change in synaptic vesicle fusion mode.

We investigated these possibilities with experiments and simulations. We constructed Monte Carlo simulations of neurotransmitter release, diffusion, and activation of kinetic models of AMPA and NMDA receptors. Simulations either changed numbers of vesicles fusing at active zones, altered the position of fusing vesicles with respect to postsynaptic receptors, or fusion pore timing. All these affected postsynaptic simulations of AMPA receptor activation. However, missalignment of fusion and receptors showed only small differential effects on AMPA and NMDA responses. MVR can govern cleft glutamate concentrations. This has been shown by investigating effects of repetitive stimulation on P_r_ (Foster et al., 2005; Christie and Jahr, 2006). Experimentally, we saw no change in glutamate cleft concentrations mediated by paired pulse stimulation to enhance P_r_, or by reducing P_r_ with raised Mg^2+^ or baclofen. This implies that putative effects of 5-HT_1B_ receptors on MVR would be more complex than a change in P_r_. In simulations we found that altering the number of vesicles fusing caused little differentiation between AMPA and NMDA receptors. This contrasts with experimental effects of 5-HT_1B_, but not GABA_B_ receptors. Our simulations favored 5-HT_1B_ receptor-mediated effects on fusion pore dilation and duration and on P_r_ for GABA_B_ receptors. Although the simulation does not account for all properties of biological fusion pores, such as charges lining a pore, or flickering (Chang et al., 2017), it is notable that sustaining a 0.4 nm pore best simulated experimental results. Biophysical data predict a similar fusion pore diameter before full fusion (Vardjan et al., 2013).

This conclusion is supported by experimental and simulated effects on response kinetics. Activation and decay rates following simulated arrested fusion pores were consistent with effects of 5-HT_1B_ receptors which similarly slowed EPSCs. In contrast, simulating changes in P_r_, at multivesicular or between univesicular synapses, sped up the decay rate or left it unchanged, while baclofen had no effect on ESPC kinetics. By targeting the release machinery, 5-HT_1B_ receptors selectively change postsynaptic receptor activation. They are ineffective inhibitors of NMDA EPSCs. This is an effect previously ascribed to kiss-and-run fusion (Choi et al., 2003; Photowala et al., 2006; Schwartz et al., 2007; Gerachshenko et al., 2009).

High Ca^2+^ concentrations allow synaptotagmin to competitively prevent Gβγ-SNARE complex binding, causing Ca^2+^-dependent loss of Gβγ-mediated presynaptic inhibition in lamprey (Yoon et al., 2007). Presynaptic Ca^2+^ accumulates during repetitive stimulation (Charlton et al., 1982; Swandulla et al., 1991). If 5-HT_1B_ receptors act by competing with Ca^2+^-synaptotagmin SNARE interactions, whether syt1 or a fusogenic protein like synaptotagmin 7 (Jackman et al., 2016), we predict loss of 5-HT-mediated inhibition during repetitive stimulation. Indeed, we demonstrate that 5-HT_1B_ receptor-mediated inhibition is prevented at high Ca^2+^ concentrations and during repetitive stimuli. This suggests that 5-HT_1B_ receptors filter synaptic transmission with an outcome similar to a silent synapse (Liao et al., 1995). This prevents stray single action potential AMPA transmission. However, high frequency stimuli, such as theta bursts, transmit efficiently.

## Conclusions

The sensitivity of 5-HT_1B_ receptor effects to Ca^2+^ accumulation and their lack of effect on NMDA responses may underly CP93129 actions on repetitive responses seen during current clamp recordings of subicular neurons. 5-HT_1B_ receptors inhibit early responses in stimulus trains, but train stimulation allows recovery. Because NMDA receptor-mediated responses are much less inhibited, they may be recruited throughout burst evoked depolarizations, even after the first stimulation when NMDA receptor-mediated responses are blocked by Mg^2+^, but the receptor is activated. Because NMDA responses are slow (Dale and Roberts, 1985; Lester et al., 1990), NMDA receptors are available for gating immediately following frequency-dependent release of inhibition of AMPA EPSPs and depolarization even during 5-HT_1B_ receptor activation.

We show that 5-HT_1B_ receptors change the profile of transmission of synaptic trains. They allow train stimuli to be transmitted, but reject responses to single stimuli. Interestingly, 5-HT_1B_ knockout mice display compromised working memory (Buhot et al., 2003), and reduced behavioral inhibition to novel stimuli, and learning is compromised in SNAP-25Δ3 mice lacking Gβγ SNARE interactions (Zurawski et al., 2019). This receptor may play a role in behaviors requiring impulse control; for example 5-HT_1B_ receptor knock-out animals are more motivated to self-administer cocaine (Rocha et al., 1998). Perhaps 5-HT_1B_ receptors selectively process information in the face of competing stimuli.

In contrast, GABA_B_ receptors inhibit Ca^2+^ entry. This implies interaction between effects of GABA_B_ and 5-HT_1B_ receptors, because the effect of 5-HT_1B_ receptor activation is modified by Ca^2+^-syt1. Indeed, low doses of baclofen modify temporal effects of CP93129 to prevent train-dependent recovery from CP9129’s inhibition, showing that receptor coactivation integrates information. Even weak GABA_B_ receptor activation prevents train dependent recovery from 5-HT_1B_ receptor activation. Thus, colocalization of presynaptic receptors responding to different agonists but whose signaling pathways converge and interact allows presynaptic terminals to act as sites of neural integration. This complex interaction also points to the need to understand relationships between Ca^2+^, release and modulation, and physiological Ca^2+^ concentrations may be lower than in this study.

## Acknowledgements

We would like to thank Drs Heidi Hamm, Janet Richmond and Jonathon Art for helpful discussions throughout the course of this study and of this manuscript. This work was funded by NIH grants R01 MH084874 and R01 NS052699 to SA, R01 MH086507 to KYT, and F31 NS063662 to EH.

---

**Multimedia Video 1:**
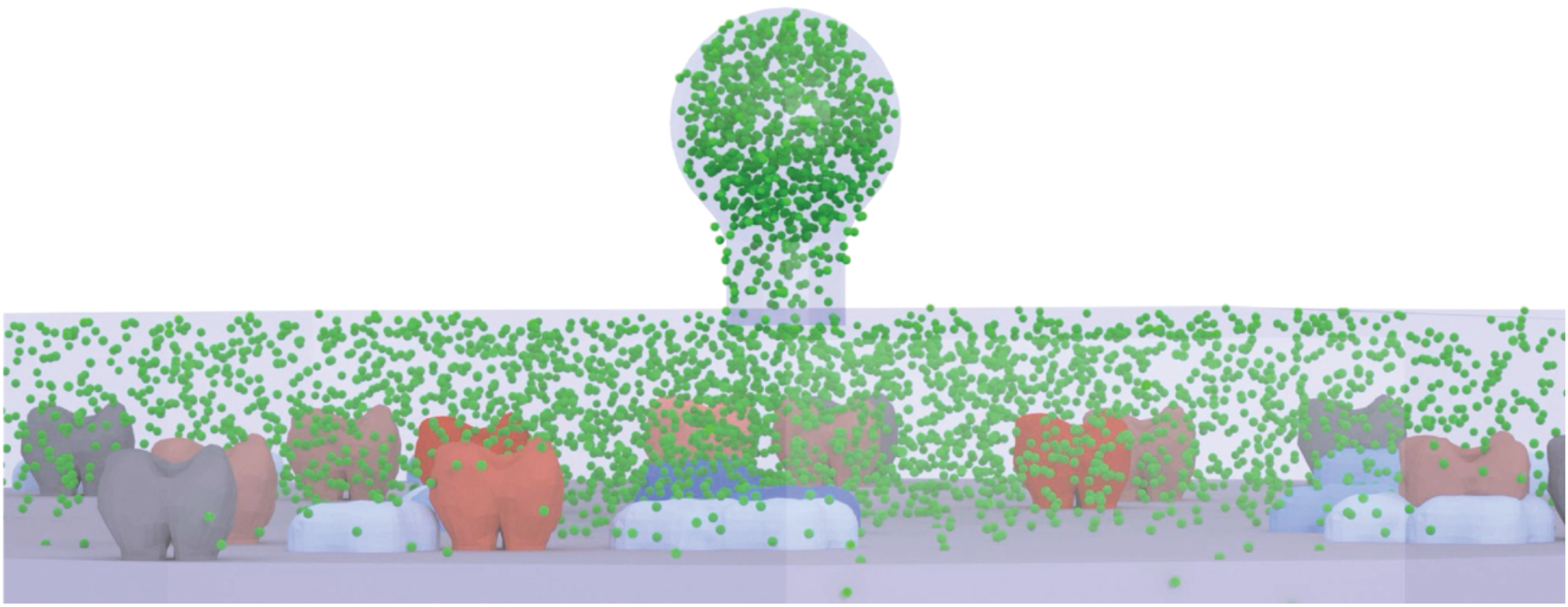
Simulation of vesicle pore opening an expansion. This video shows the simulated synapse from the side demonstrating fusion of the vesicle compartment with the synaptic cleft, release of glutamate molecules (green) and binding of glutamate to post synaptic AMPA receptors (red shades) and NMDA receptors (blue shades). Receptor colors depend on kinetic state (see Fig 7D for kinetic models and colors).

**Multimedia Video 2:**
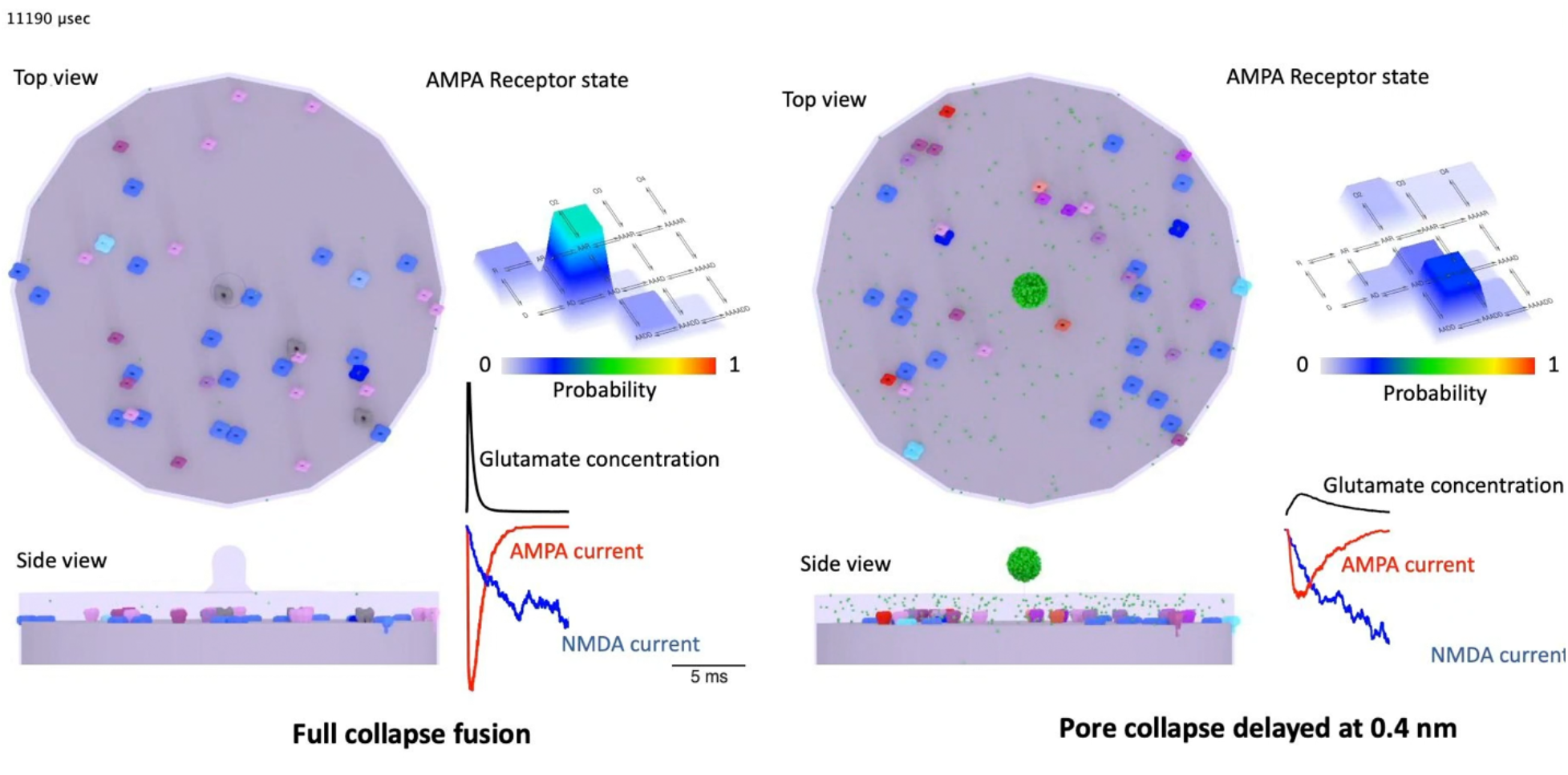
Simulations of full rapid fusion and delayed fusion pore opening. Left hand panels – top and side views of vesicle fusing and releasing glutamate into the synaptic cleft. AMPA receptor state probabilities are shown as a color coded lookup in 3D relief. Simulated cleft glutamate concentrations (black), AMPA receptor current (red) and NMDA receptor current (blue) are shown up to the clip time point. Right hand panels show the same but during delayed fusion pore opening. Receptor colors are dependent on state (Fig 7D). The still is of the video 11.19 ms after the start of fusion.

## Notes

Conflict of Interest: none

https://alford.lab.uic.edu/GPCRs.html

### Competing Interest Statement

The authors have declared no competing interest.

### Summary of Updates

Updates to demonstrate the difference in effects of GABAB and 5-HT1B receptor activation on glutamate release detected by iGluSnFR were made to Figure 6. Additionally, point spread functions of the lattice light sheet microscope were provided.

